# Proteome Analysis of Soybean Root Apoplast Combined with AlphaFold Prediction Reveal *Macrophomina phaseolina* Infection Strategies and Potential Targets for Engineering Resistance

**DOI:** 10.1101/2024.12.29.630648

**Authors:** Chetan Veeraganti Naveen Prakash, Muthusaravanan Sivaramakrishnan, Sakshi Goel, Vasudha Porwal, Pawan Kumar Amrate, Manoj Kumar Shrivastava, Balakumaran Chandrasekar

## Abstract

*Macrophomina phaseolina* (Tassi) Goid. is a hemibiotrophic pathogen that causes charcoal rot (CR) disease in various legumes, including soybean. To date, no reliable resistance gene sources have been identified in soybean or other legumes to combat *M. phaseolina*. Therefore, the identification of mechanistic targets is crucial for improving resistance against the pathogen. The apoplast is a critical region where intense molecular cross-talk occurs between plants and pathogens, and the outcome of their interactions is determined in this compartment. Here, we employed label-free quantitative (LFQ) proteomics to investigate the dynamics of soybean root apoplast during *M. phaseolina* infection. We have detected several secreted proteins of *M. phaseolina* and differential regulation of soybean-secreted proteins in root apoplast during infections. Glycome analysis and callose deposition assays have revealed changes in soybean root cell wall compositions and potential polysaccharide targets of *M. phaseolina*. AlphaFold 2 (AF2) analysis was instrumental in revealing several interesting sequence-unrelated structurally similar (SUSS) effectors and effectors with novel structural folds secreted by *M. phaseolina*. Structured-guided engineering of protease-inhibitor complexes is emerging as an important strategy to engineer resistance in plants against pathogens. AlphaFold Multimer (AFM) analysis of candidate-secreted proteins from soybean and *M. phaseolina* has predicted cysteine and serine protease-inhibitor complexes with high confidence. We have validated these interactions using molecular dynamics (MD) and competitive activity-based protein profiling (ABPP) approaches. Therefore, our work provides insights into Soybean-*M. phaseolina* interactions in the root apoplast and unveil potential candidates for engineering resistance.

## Introduction

*Macrophomina phaseolina* (Tassi) Goid. is a devastating soil-borne fungal pathogen that causes charcoal rot (CR) disease or dry root rot (DRR) disease in several crops under high temperatures and drought stress conditions. *M. phaseolina* has been predicted to pose a serious threat to several legumes under changing climatic conditions (Pandey and Basandrai 2021). Among their various legume hosts, soybean (*Glycine max* L. Merril) is one of the most economically important crops that are vulnerable to CR disease with yield losses ranging from 70-100% (Amrate et al. 2023; Ortiz et al. 2023). During root colonization, *M. phaseolina* exhibits an initial mycelial phase followed by the microsclerotia stage, leading to characteristic charcoal rot disease symptoms (Chowdhury et al. 2017). The microsclerotia phase of the pathogen plays a critical role in its pathogenesis. Therefore, it is crucial to identify suitable candidates in crop plants for engineering resistance against the pathogen.

Apoplast is an important compartment where the initial dialogue between the plant and pathogen occurs and the dynamics of apoplastic components determine the outcome of plant-pathogen interaction. All the components outside the cell membrane, including the cell wall, constitute the apoplast. The cell wall is a dynamic component composed of various cellulosic and hemicellulosic polysaccharides that serve as a protective barrier to resist pathogen invasion. Furthermore, at the pathogen infection sites, the cell wall is reinforced with appositions such as callose deposition that act as barriers to pathogen invasion (Hosseini-Zahani and Taheri 2023). Fungal pathogens employ several strategies for the successful colonization of their host plants (Buscaill and van der Hoorn 2021). They secrete carbohydrates-active enzymes (CAZymes) that are involved in plant cell wall degradation to gain entry into the host (Sivaramakrishnan et al. 2024). They also manipulate several host-derived factors that act as susceptible factors to favor pathogen colonization (Hong et al. 2024). Furthermore, fungi secrete effectors in apoplast to suppress host immune responses. Some effectors play a role in hijacking host defense machinery by inducing cell death following the inverse gene for gene interactions (Di et al. 2024). Compared to the biotrophic fungal pathogens, only a few necrotrophic effectors have been identified, and their biology is largely understudied (Shao et al. 2021). A large number of necrotrophic effectors are yet to be identified, and machine learning tools such as AlphaFold (AF) would assist in the discovery of novel effectors, which would aid in formulating efficient strategies to control the fungal pathogen (Seong and Krasileva 2023).

Protease-inhibitor interaction is another important aspect that occurs in the apoplast during fungal colonization and determines the outcome of the plant-pathogen interaction (Misas-Villamil et al. 2016). Plant-secreted proteases such as cysteine and serine proteases in the apoplast represent central hubs for plant immune responses (Godson and van der Hoorn 2021). These enzymes mediate the generation of immunogenic elicitors or directly cleave the proteins essential for pathogen virulence and thereby impart resistance in plants (Paulus et al. 2020; S. Wang et al. 2021). The activities of these immune proteases are regulated by the inhibitors that are produced by the pathogen or the plant itself (Wang et al. 2020). AlphaFold multimer (AFM) analysis is emerging as an important tool in plant biology for structural-level understanding and prediction of protein-inhibitor interactions at the plant-pathogen interface (Homma et al. 2023; Mooney and van der Hoorn 2024). AFM analysis has been instrumental in designing strategies for modulating apoplastic protease-inhibitor interactions for enhancing resistance against the oomycete pathogen *Phytophthora infestans* (Schuster et al. 2024). So far, reliable resistance gene sources have neither been identified in soybean nor in other legumes to combat *M. phaseolina*. Therefore, predicting apoplastic protease-inhibitor interactions would assist in formulating strategies for engineering resistance in these crops against the pathogen.

Proteomics is an important approach to investigate the dynamics of apoplast during pathogen infections. To date, the apoplastic proteome studies are largely focused on foliar pathogens, and the studies on root-invading fungal pathogens are limited. This is due to the fact that a large number of plants are needed to collect sufficient apoplastic fluids from roots for proteome analysis. Notably, the apoplastic proteome analysis of soybean roots and their dynamics during *M. phaseolina* has not been investigated. Here, in our study we have performed label-free quantitative proteomics of the apoplastic fluids collected from soybean roots during *M. phaseolina* infection. In our proteome analysis, we detected several secreted proteins of *M. phaseolina* and differential regulation of several soybeans secreted proteins in root apoplast during infections. Glycome analysis and callose deposition assays have revealed changes in soybean root cell wall compositions and potential polysaccharide targets of *M. phaseolina*. AlphaFold 2 (AF2) analysis was instrumental in revealing several interesting sequence-unrelated structurally similar (SUSS) effectors and effectors with novel structural folds secreted by *M. phaseolina*. Structured-guided engineering of protease-inhibitor complexes is emerging as an important strategy to engineer resistance in plants against pathogens. AlphaFold Multimer (AFM) analysis of candidate-secreted proteins from soybean and *M. phaseolina* has predicted cysteine and serine protease-inhibitor complexes with high confidence. We have validated these interactions using molecular dynamics (MD) and competitive activity-based protein profiling (ABPP) approaches. Therefore, our work provides insights into Soybean-*M. phaseolina* interactions in the root apoplast and unveil potential candidates for engineering resistance.

## Material and methods

### Isolation and identification of *Macrophomina phaseolina*

Infected root samples were collected from Jawaharlal Nehru Krishi Vishwa Vidyalaya (JNKVV), Jabalpur, Madhya Pradesh, India (23° 12’ 47.808’’ N, 79° 57’ 49.752’’ E, 301.5 m above mean sea level). Root bits were surface sterilized using 0.1% mercuric chloride and rinsed thrice with sterile Milli Q water. The surface sterilized bits were placed on Potato Dextrose Agar and incubated at 28℃. Hyphal tip method was followed to obtain pure cultures of the pathogen (Brown 1924). Molecular identification of the fungus was performed by Sanger sequencing of Internal Transcribed Spacer 1/4 (ITS 1/4) (Primers: ITS1: TCCGTAGGTGAACCTGCGG, ITS4: TCCTCCGCTTATTGATATGC). The obtained consensus sequences of forward and reverse sequences from Sangar sequencing were analyzed in NCBI-BLAST. The phylogenetic tree was constructed with the neighbor-joining method using MEGA 11 software. The Maximum Composite Likelihood model was selected and branch support was calculated by 1,000 Bootstrap repetitions. The ITS1/4 sequence was submitted to NCBI Genbank with accession no. PQ807640.

### Plant material

The soybean seeds of varieties JS 20-94, JS 20-34 and JS 20-98 were procured from Jawaharlal Nehru Krishi Vishwa Vidyalaya (JNKVV), Jabalpur, Madhya Pradesh, India and SL-955 was procured from Department of Plant Breeding and Genetics, Punjab Agricultural University, Ludhiana, Punjab, India. These varieties were used for the pathogenicity test of the isolated *M. phaseolina* strain.

### Root dip method of infection with *M. phaseolina* culture

Infection assay was performed as described by Nelson et al. 2021 with slight modifications. Briefly, the healthy soybean seeds were sterilised by chlorination for 16 hours (Sivaramakrishnan et al. 2023). Sterilized seeds were sown in double autoclaved soil mixture (Soilrite) and were grown in a plant growth chamber (28℃, 75% humidity, 16:8 hour day: night schedule, 6000 lux luminescence) for 2 weeks. These plants were washed in tap water and dipped in 25% inoculum (w/v) of actively growing mycelium of *M. phaseolina* (*M. phaseolina_Jbstrain1)* cultured in Potato Dextrose Broth. Milli Q water was used as control. Later, the plants were repotted and placed in a plant growth chamber (35℃, 65% humidity, 16:8-hour day: night schedule, 6000 lux luminescence). Plants were watered at 8-hour intervals and the disease progression was recorded for 3 days.

### Microscopic analysis of soybean roots upon *M. phaseolina* infection

Control and infected soybean roots were prepared for microscopic analysis as described by Barrow (2004) with slight modifications. The roots collected at different stages of *M. phaseolina* infection were subjected to overnight clearing of roots with 70% ethanol. The cleared roots were boiled in 10% KOH at 100℃ for 1 hour. The roots were washed with tap water, bleached with 10% alkaline H_2_0_2_ for 45 minutes and acidified with 1% HCl for 2 minutes. These cleared roots were mounted on a glass slide using acidified glycerol. Differential Interference Contrast (DIC) imaging was performed using Fully Spectral Confocal Laser Scanning Microscope (FSCLSM, Carl Zeiss LSM 880).

### Root apoplastic fluid extraction

The root apoplastic fluids were collected employing vacuum infiltration and centrifugation method (Chandrasekar et al. 2014, Chandrasekar et al. 2022). In brief, the washed and cut pieces of soybean roots were submerged in ice-cold Milli Q water, and a vacuum was applied. The water infiltrated into the intercellular spaces of roots as the vacuum was slowly released. The roots were then dried briefly and centrifuged at 1000 g and 4°C for 30 min using a swing bucket rotor. Around 45 control plants and 90 *M. phaseolina* infected plants were used to collect approximately 3 ml of apoplastic fluid. The apoplastic fluid collected was used for proteome analysis and ABPP experiments.

### Label-free quantitative proteomics of root apoplast using LC-MS/MS

Apoplastic fluid was reduced with 5 mM TCEP, alkylated with 50 mM iodoacetamide, and then digested with Trypsin (1:50, Trypsin/lysate ratio) for 16 h at 37 °C. Digests were cleaned using a C18 silica cartridge to remove the salt and dried using a speed vac. The dried pellet was resuspended in buffer A (2% acetonitrile, 0.1% formic acid). Experiments were performed on an Easy-nlc-1000 system coupled with an Orbitrap Exploris mass spectrometer. 1ug of peptide sample were loaded on C18 column of 15 cm length, 3.0μm Acclaim PepMap (Thermo Fisher Scientific) and separated with a 0–40% gradient of buffer B (80% acetonitrile, 0.1% formic acid) at a flow rate of 500 nl/min and injected for MS analysis. LC gradients were run for 110 minutes. MS1 spectra were acquired in the Orbitrap (Max IT= 60ms, AGQ target= 300%; RF Lens= 70%; R= 60K, mass range= 375−1500 m/z; Profile data). Dynamic exclusion was employed for 30s excluding all charge states for a given precursor. MS2 spectra were collected for top 20 peptides (Max IT= 60ms, R= 15K, AGC target 100%). The mass spectrometer was set in the positive ion mode. The mass spectrometry proteomics data have been deposited to the ProteomeXchange Consortium via the PRIDE [1] partner repository with the dataset identifier PXD059244.

### Proteome data analysis

The mass spectrometer was operated using Proteome Discoverer (v2.5) software. For dual Sequest and amanda search, the precursor and fragment mass tolerances were set at 10 ppm and 0.02 Da, respectively. Both peptide spectrum match and protein false discovery rate were set to 0.01 FDR. The obtained proteome data was analyzed to predict the putative secretory proteins using Signal P 6.0. The signal P domains containing proteins were then identified and classified as CAZymes using dbCAN3 server (last accessed: June, 2024) using HMMER dbCAN (E-value < 10−15, coverage > 0.35), DIAMOND (E-value < 10−102), and HMMER dbCAN-sub (E-value < 10−15, coverage > 0.35) tools (Zhang et al., 2018). Only the proteins successfully annotated by at least two of the tools were considered CAZymes. The rest of the proteins were searched for their domain architecture under Pfam which is now hosted by InterPro (Mistry et al. 2021) (last accessed: July, 2024). The secreted proteins were subjected to EffectorP Fungi 3.0 algorithm for the prediction of effector candidates (Sperschneider and Dodds 2022). Volcano plots were generated using EnhancedVolcano package, the PCA was performed using factoextra package and Pearson’s correlation was visualised using corrplot package in R software. Z-score values of the differentially regulated plant proteins were calculated using the formula z= (X-µ)/σ where X= individual abundance value (LFQ), µ= mean of the replicate abundance values (LFQ), σ= Standard deviation of the replicate abundance values (LFQ) and was illustrated as a heat map using TBtools-II v1.112. The sequences of functionally characterized AA9s were collected from UniProt database (The UniProt Consortium 2024) and a phylogenetic tree was constructed along with the detected members of AA9 with the neighbor-joining method in MEGA 11 software.

### GO annotation of the proteome dataset

Gene Ontology (GO) analysis of the differentially regulated soybean apoplastic proteins based on molecular functions and biological processes was performed using g:GOSt module of g:Profiler with Fisher one-tailed test and Benjamini-Hochberg with an FDR threshold of ≤0.05 to correct for multiple testing. GO analysis fungal-secreted proteins in terms of molecular functions and biological processes was performed using FungiFun v2.2 with hypergeometric distribution test and Benjamini-Hochberg FDR threshold of ≤ 0.05.

### FTIR spectroscopy of root AIR samples

FTIR analysis was performed using Fourier Transform Infrared Spectroscopy (Model: Bruker Alpha) over a range of 4,000 to 400 cm^−1^. For each spectrum, 32 scans were performed at a resolution of 4 cm^−1^. The spectra were baseline-corrected and smoothened using OPUS spectral processing software with default parameters. The functional groups were annotated based on corresponding wavenumbers.

### Monosaccharide composition and glycosyl linkage analysis of root AIR samples

To determine the monosaccharide composition of the control and infected root samples, alditol acetate derivatization was performed (Foster et al. 2010; Pettolino et al. 2012). Initially, alcohol insoluble residue (AIR) samples of the roots were obtained by washing sequentially with 70% ethanol, chloroform:methanol (1:1), and acetone. Then, the AIR samples were hydrolyzed with 2.5 M trifluoroacetic acid (TFA), reduced with sodium borohydride (NaBH_4_) in NH_4_OH followed by acetylation. The alditol acetates were extracted using ethyl acetate and water by phase-separation centrifugation. The organic phase was collected, dried under N_2_ and resuspended in acetone. The derivatised sugars were analyzed by GC-MS on a 30 m × 0.25 mm SP-2380 Fused Silica Capillary Column (Supelco). Injection was at 80°C with a 3-min hold, ramped to 170°C at 30°C min^−1^, and then to 240°C at 4°C min^−1^, with a 20-min hold at the upper temperature. Helium flow was 1.5 mL min^−1^ with split-less injection. The electron impact mass spectrometry was performed with model no. TQ8040 CI/NCI (Shimadzu) at 150 eV and a source temperature of 250°C. The obtained chromatograms were annotated based on retention time and the mass spectrum of standard sugars. Glycosidic linkage analysis was done as described in (Chandrasekar et al. 2022; Kim et al. 2020; Liu et al. 2015). Briefly, dry DMSO was added to AIR samples and kept for stirring overnight at room temperature. NaOH/DMSO was added, and methylation was carried out by adding methyl iodide under a Nitrogen environment for 4 hours with stirring. The reaction was quenched with Milli Q water, bubbled with nitrogen, and dichloromethane (DCM) was added. The DCM phase was collected, dried, and subjected to hydrolysis and alditol acetate derivatization as followed in monosaccharide composition analysis to obtain partially methylated alditol acetates. These derivatised sugars were analyzed using similar parameters as that of monosaccharide composition analysis with GC-MS. The glycosidic linkages were assigned based on retention time and mass spectrum fragmentation patterns compared to the CCRC spectral database (https://www.ccrc.uga.edu/specdb/ms/pmaa/pframe.html).

### Crystalline cellulose estimation

Crystalline cellulose estimation of AIR samples of control and infected roots was performed as described by (Pettolino et al. 2012). The pellet after the TFA hydrolysis was treated with Updegraff reagent followed by Saeman hydrolysis. The samples were subjected to anthrone assay to estimate the released glucose, and the readings were taken using a microtiter plate reader at 625nm. Absolute quantification of the crystalline cellulose was calculated against a standard curve plotted using various concentrations of glucose standards from 0-10 µg/ml.

### Root callose staining and its quantification

Callose staining of the soybean roots was performed using 0.01% Aniline Blue as described by Schenk and Schikora 2015 with few modifications. In brief, the soybean roots collected at different stages of *M. phaseolina* infection were washed with Milli Q water to remove soil particles and kept in acetic acid: ethanol (1:3) overnight. The roots were transferred to 10% KOH solution and were heated in a water bath at 95℃ for 2 minutes. Subsequently, they were washed with Milli Q water and 1X PBS for 30 minutes each thrice. Then, the roots were washed with 150mM K_2_HPO_4_ for 1 hour, followed by staining with 0.01% Aniline blue in 150mM K_2_HPO_4_ for 1 hour and again washed with 150mM K_2_HPO_4_ for 1 hour. The stained roots were visualized under Zeiss ApoTome 2.0 microscope (Carl Zeiss) using a DAPI filter with an excitation wavelength of 370 nm and an emission wavelength of 509 nm (magnification= 10X). The callose quantification was performed as previously described (Mason et al. 2020). The images were scaled and pre-processed using the “Clear Measure Resize” macro to set the background colour to black, deleted pixels outside the selected area, measured the properties of the selected area, and resized the image to specific dimensions using bilinear interpolation. The resulting images were segmented to detect callose deposits using the Trainable Weka Segmentation plugin of ImageJ with default parameters. The segmented image was converted to an 8-bit format with the background set to black, transformed into a mask, and analyzed for particles of a size (0.000005–0.005), circularity parameters (0.25–1.0), and counted (Arganda-Carreras et al. 2017; Schindelin et al. 2012). Callose deposits were expressed as the number of deposits per area.

### SUSS effector identification using AlphaFold2

The protein structures were predicted using AF2 as a monomer (Jumper et al. 2021; Mirdita et al. 2022). To refine the complex structure, relaxation was performed using the amber force field. The predicted AF models were compared to the experimentally resolved PDB100 database using Foldseek search in TM-align mode with taxonomic restriction to Ascomycota (van Kempen et al. 2024). All TM-scores were normalized relative to the length of the experimentally resolved proteins and TM-score >0.5 were considered as structural analogs. The superposition of predicted AF models onto the identified PDB hits was performed using the Matchmaker module of UCSF chimera (Meng et al. 2006; Pettersen et al. 2004). The 2D topology maps for the modelled SUSS effectors were predicted using the PDBsum webserver (Laskowski et al. 2017).

### AlphaFold Multimer (AFM) analysis

A high-throughput *in silico* pulldown screening with AlphaFold v2.3.2 implemented in the LocalColabFold v1.5.5 (https://github.com/YoshitakaMo/localcolabfold), a local installation of ColabFold was performed. All complexes are predicted as heterodimers using LocalColabFold v1.5.5 with the AF multimer model using templates from the MMseqs2 (combined paired and unpaired MSA) search. The top 20 hits with an E-value of < 0.1 were selected as candidate templates for multiple sequence alignment (MSA) and subsequent input parameters for AFM. We ran the AFM model on the GPU with the following parameter settings: template usage enabled, 20 recycles, early recycle stop tolerance set to 0.5 to control the convergence criterion, no dropout, and energy minimized using amber force field. All runs were performed on a Linux system with an Intel CORE i9 11th generation processor, 32 GB RAM and 6 GB NVIDIA RTX 3060 GPU. We evaluated the predicted protein pairs based on AlphaFold2 performance metrics such as predicted template modelling (pTM) score, predicted local distance difference test (pLDDT), and interface pTM score (ipTM). The above performance metrics were parsed using the in-house Python script (https://github.com/chemicalglycobiologylab) from LocalColabfold .json output files. We also calculated spatial information about interface residues as well as confidence and accuracy metrics at the residue and contact levels using the Colabfold Batch AlphaFold-2 multimer structure analysis pipeline to determine whether a pair of residues is a valid contact, particularly at the heterodimer interface and the catalytic triad (Schmid 2023). We found experimentally resolved protein structures from the PDB with structural folds similar to predicted heterodimers using Foldseek multimer (van Kempen et al. 2024). All TM scores were adjusted based on the query length and experimentally solved template structures. The structural similarities of modelled proteins were assessed using the DALI webserver (Holm 2022). The interface of the predicted heterodimer complexes was analyzed and visualized using the open-source version of the PyMOL Molecular Viewer (Yuan, Chan, and Hu 2017), and contact maps were created using the PDB module of Biopython with a contact cut-off of 8 Å (Cock et al. 2009).

### Molecular dynamics simulation studies

The protein-protein complexes are simulated for 100ns using the OPLS-AA force field (Lindahl et al. 2010) implemented in GROMACS 2022.3 (Lemkul 2019). All the structures were equilibrated using Constant temperature, constant volume (NVT) and Constant temperature, constant pressure (NPT) ensembles for a constant temperature of 303 K and 1 atm pressure. Necessary amounts of sodium (Na) and chloride (Cl) were added to the system for neutralization. Periodic boundary conditions were enforced in all directions. All the bond lengths within the system were constrained using the LINear Constraint Solver (LINCS) algorithm. Long-range electrostatics in the system was computed by using the Particle Mesh Ewald method (PME). The V-rescale weak coupling method was used to implement temperature regulation at 310 K. The NVT and NPT ensembles were both equilibrated using the Parrinello-Rahman coupling method. Once the system was equilibrated with desired temperature and pressure, the last stage production run was performed for 100 ns. The frames were saved at 10-ps intervals throughout the MD run, with a time step of 2 fs. The built-in modules of GROningen MAchine for Chemical Simulations (GROMACS), such as gmx_rms, gmx_rmsf, gmx_dist and gmx_hbond, were used to evaluate root mean square deviation (RMSD), root mean square fluctuation (RMSF), pairwise distance between chains, and the number of hydrogen bonds (Hbond) along the entire MD trajectory. The structures were visualized using the PyMOL and Chimera software (Meng et al. 2006; Yuan et al. 2017). The Free energy landscape (FEL) analysis was performed using gmx_anaeig module and free energy landscape v1.0.3 module available at https://github.com/sulfierry/free_energy_landscape. The binding free energy was determined by utilizing the 100 ns trajectory obtained from molecular dynamics simulations, and calculations were performed on every 100 frames extracted from the last 30 ns of the trajectory using the gmx_mmpbsa and hawkdock (Valdés-Tresanco et al. 2021; Weng et al. 2019).

### Competitive activity-based protein profiling (ABPP) of apoplastic fluids and fungal culture filtrate with soybean Kunitz

Competitive ABPP was performed as described by (Chandrasekar et al. 2017; Hong and Hoorn 2014; Kaschani et al. 2012). In brief, apoplastic fluids collected from control and infected soybean roots at 3 days post infection (dpi) were preincubated with commercially available soybean Kunitz (Sigma Cat no. 93620-250MG) at concentrations 1.25, 2.5, 5, and 7.5 µg/ mL for 30 minutes. Milli Q was used as control. The apoplastic fluids were treated with the FP-alkyne probes or DMSO (no-probe control) along with 500mM Sodium acetate of pH 6 to maintain the pH of the reaction. The mixtures were kept on a thermoshaker (37℃, 200 rpm, 2 hours). The labeling reactions were quenched by ice-cold acetone precipitation. The protein pellets were resuspended in 1X PBS-SDS buffer, heated at 90°C for 10 min, and used for click chemistry. For click-chemistry, 1mM N3Rh (Final conc. 50µM), 3.4mM TBTA (Final conc. 100µM), 100mM TCEP (Final conc. 2mM), and 50mM CuSO_4_ (Final conc. 1mM) was added to the sample and was kept on a thermoshaker under dark conditions (30℃, 800 rpm, 1 hour). The reactions were quenched by acetone precipitation and the protein pellets were dissolved in 1X PBS-SDS buffer and 4X gel loading buffer (Final conc. 1X). The labelled samples were separated on 10% SDS-PAGE gels at 200V for 45 minutes and fluorescently labelled proteins were detected by in-gel fluorescence using a ChemiDoc MP scanner 532/28 excitation/emission filter of Pro-Q Emerald 488 in the Blue Epi filter. Densitometric analysis of the fluorescent bands was performed using Image Lab 6.1.0 software. Competitive ABPP was performed on the culture filtrate that was collected from the 3-day-old liquid cultures inoculated with *M. phaseolina*. The culture filtrate was filtered using Mira cloth and was preincubated with various concentrations of soybean Kunitz. The preincubated culture filtrate was subjected to the above-mentioned ABPP protocol.

## Results

### *Macrophomina phaseolina* secretes a large repertoire of CAZymes in the soybean root apoplast

Here, first, we isolated the native *Macrophomina phaseolina* strain from the soybean fields of Jabalpur, India, a hot spot region of charcoal rot (CR) disease. The pure cultures of *M. phaseolina* were isolated from the infected soybean roots using the hyphal tip method (Figure S1A). To confirm the identity of the isolated strain, the Internal Transcribed Spacer (ITS) was amplified and sequenced (Figure S1B). Basic Local Alignment Search Tool (BLAST) analysis against the National Center for Biotechnology Information (NCBI) database has revealed a 100% match with the *M. phaseolina* strains that have been previously isolated (Figure S1C). To test the pathogenicity of the isolated *M. phaseolina* strain (*M. phaseolina_Jbstrain1*), the roots of two-week-old seedlings of elite Indian soybean varieties such as JS 20-94, SL 955, JS 20-34 and JS 20-98 were infected using the root dip inoculated method. The characteristic above-ground CR disease symptoms such as leaf chlorosis, marginal necrosis, and wilting were prominently observed during the 3 days post-infection (dpi) (Figure S2). Of the tested soybean varieties, JS 20-94 displayed severe above-ground symptoms (Figure S2). These results indicate that the isolated native *M. phaseolina* strain is pathogenic on various soybean varieties, and the soybean variety JS-2094 is highly susceptible to *M. phaseolina*. Therefore, we used the JS-2094 soybean variety and the isolated virulent *M. phaseolina* strain for our further studies.

To monitor the colonization process of isolated *M. phaseolina* strain, the roots of two-week-old seedlings of the JS 20-94 soybean variety were infected with the mycelia of *M. phaseolina* using the root-dip inoculation method. The root samples were collected at three different time points: 1 dpi, 2 dpi, and 3 dpi. The differential Interface Contrast (DIC) microscopic analysis of the collected samples has revealed the formation of microsclerotia structures by the pathogen during the process of colonization (Figure 1A). At 3 dpi, the number of microsclerotia was higher compared to earlier time points (Figure 1A). To investigate the proteins that are secreted by *M. phaseolina* and monitor the dynamics of secreted soybean proteins in the apoplast, the roots of two-week-old seedlings of the JS 20-94 soybean variety were infected with the mycelia of *M. phaseolina*. The apoplastic fluids were collected from the infected and control roots at 3 dpi and subjected to shotgun proteome analysis (Figure 1B). In total, we have detected 1543 proteins in our apoplastic proteome data set (Table S1). The Principal Component Analysis (PCA) of these detected proteins has indicated a clear distinction between the control and infected proteome samples and a positive correlation within the respective biological replicates (Figure S3). Analysis of these proteins using the SignalP server has revealed that 483 proteins (31%) carried the secretory signal peptide (SP) at their N-terminus (Figure 1B and Table S2). The remaining proteins detected in the root apoplast might be derived from noncanonically secreted proteins or cytoplasmic contaminations during extraction. Only those proteins that carried the SP were considered for further analysis. Of these 483 proteins, 228 secreted proteins were from *M. phaseolina* and 255 secreted proteins from soybean (Figure 1B). First, we analyzed the proteins secreted by *M. phaseolina* in the soybean root apoplast (Table S3). Gene ontology analysis has indicated that the *M. phaseolina* secreted proteins are associated with biological processes such as including carbohydrate metabolism, proteolysis, and cell adhesion (Figure 1C). Furthermore, this analysis has revealed that these proteins are associated with several molecular functions targeting various polysaccharides and protein substrates (Figure 1C). Pfam analysis has revealed that the majority of *M. phaseolina* secreted proteins are CAZymes (126 proteins, 55%), followed by 25 proteases, 61 other diverse proteins, and, notably, 16 proteins whose domain information is not available (Figure 1D). The detected 126 CAZymes belong to various classes, such as glycosyl hydrolases (GHs, 77 proteins), auxiliary activity family enzymes (AAs, 22 proteins), carbohydrate esterases (CEs, 15 proteins), polysaccharide lyases (PLs, 8 proteins), and other unclassified CAZyme classes (4 proteins). The 77 detected GHs belonged to 36 different GH families, of which members of the GH43 family are detected in large numbers (14 proteins) (Figure 1D). A majority of members of the GH43 family have been shown to display α-L-arabinofuranosidase and/or β-D-xylosidases activities, suggesting the polysaccharides carrying linkages might be targeted by *M. phaseolina* during infections. The auxiliary activity family enzymes (AAs) represent the CAZymes that are detected in large numbers following GHs. The 22 AAs belong to seven different families (AA1, AA2, AA3, AA5, AA7, AA8, and AA9) (Table S3). The members of AA9 are Lytic Polysaccharide Monooxygenases (LPMOs) that predominantly target recalcitrant crystalline polysaccharides of the plant cell wall. The phylogenetic tree analysis of the detected AA9 members with the characterized LPMOs from various fungi suggests that these enzymes might target various plant cell wall polysaccharides such as cellulose, xylan, xyloglucan, glucomannans and mixed linkage glucans (Figure S4). The 15 CEs belong to eight different CE families (CE1, CE2, CE4, CE5, CE8, CE12, CE15, and CE16), the eight PLs belong to four different PL families (PL1, PL3, PL4, and PL9). Taken together, these results indicate that *M. phaseolina* secretes a diverse repertoire of various CAZymes belonging to various families in the soybean apoplast, and carbohydrate metabolic processes in the soybean root apoplast might play an important role during infections.

**Figure 1:**
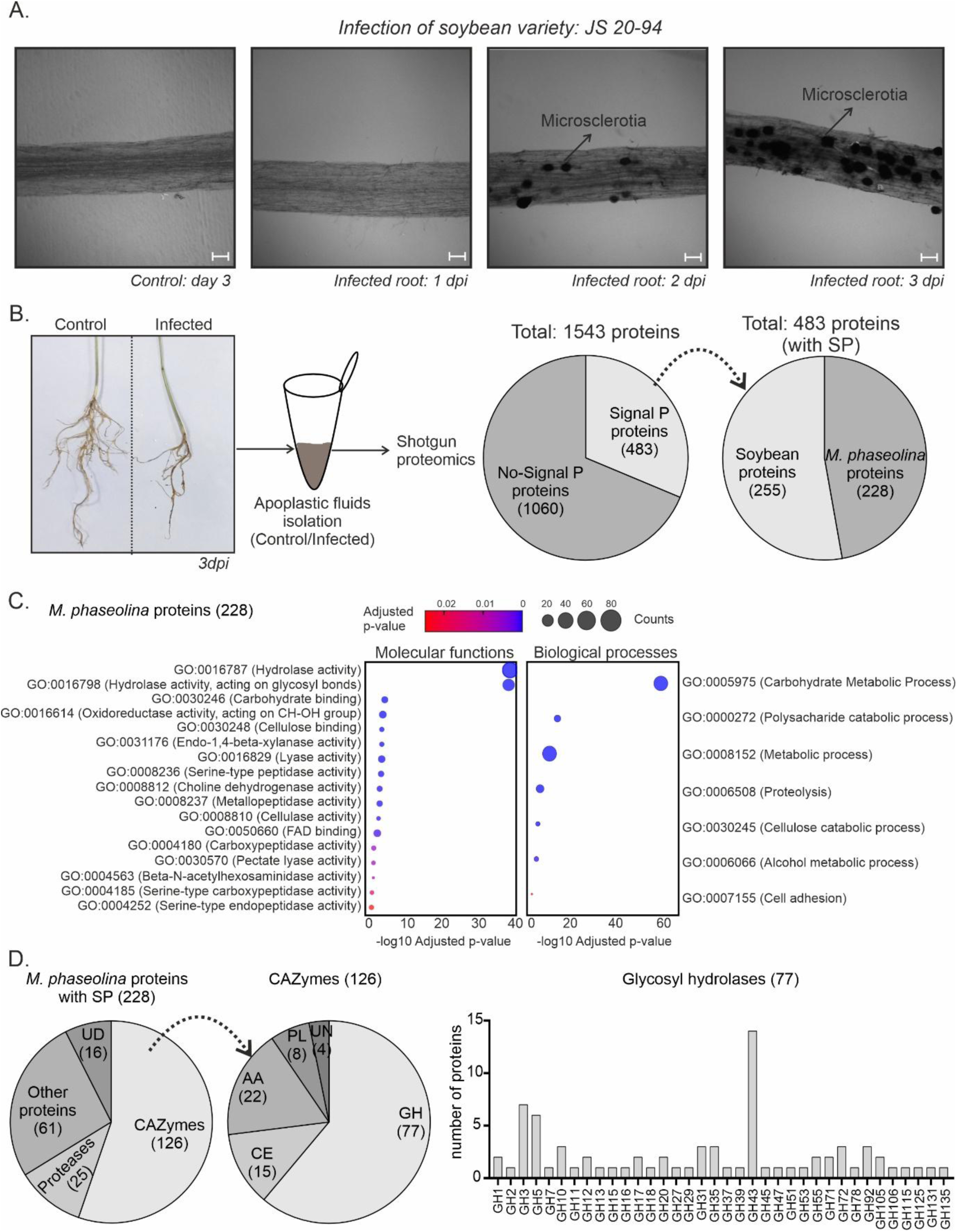
*Macrophomina phaseolina* secretes a large repertoire of carbohydrate-active enzymes (CAZymes) in the soybean root apoplast during infection. (A) Microscopy of soybean root colonization by *M. phaseolina*. The two-week-old seedlings of highly susceptible soybean variety, JS 20-94, were treated with 25% (w/v) of homogenised actively growing *M. phaseolina* culture using root-dip inoculation method. The roots collected from 1 dpi, 2 dpi and 3 dpi were washed and cleared overnight with 70% ethanol. Fungal colonisation in the soybean roots was visualised using Differential Interface Contrast (DIC) microscopic analysis (magnification= 10X, scale = 100µm). The arrow marks indicate the microsclerotia structure formed in the infected roots at 2 dpi and 3 dpi, respectively. (B) Shotgun proteome analysis of apoplastic fluids collected from control and *M. phaseolina*-infected soybean roots. Apoplastic fluids were collected from 2-week-old control and *M. phaseolina*-infected soybean plants at 3 dpi. Trypsin-digested peptides of the control and infected apoplastic fluids was subjected to shot-gun proteome analysis using LC-MS/MS. Signal P 6.0 (http://www.cbs.dtu.dk/services/ SignalP/) tool was used to identify the proteins with signal peptides (C) Gene ontology analysis of proteins secreted by *M. phaseolina*. Gene ontology analysis was performed using FungiFun v2.2 with a hypergeometric distribution test and Benjamini-Hochberg FDR threshold of ≤ 0.05. The colour of the bubbles represents the adjusted p-value denoting its statistical significance, and the size of the bubbles represents the counts, i.e., the number of genes involved in a particular molecular function or biological process. GO IDs represent taxonomy that statistically determines the involvement of those sets of genes in the specific biological processes and molecular functions as indexed in QuickGo (https://www.ebi.ac.uk/QuickGO/) (D) Classification of proteins secreted by *M. phaseolina*. The proteins were classified into CAZymes, proteases, other proteins, and unknown domain (UD) based on their domain architecture, which was predicted by Pfam domain analysis. The CAZymes were classified based on the families such as glycosyl hydrolases, carbohydrate esterases, auxiliary activities, and polysaccharide lyases using dbCAN3 server. dpi, days post-infection, SP, signal peptide, UD, unknown domain, GH, glycosyl hydrolases, PL, polysaccharide lyases, CE, carbohydrate esterases, AA, auxiliary activities, UN, unclassified CAZymes

### Cell wall composition and callose depositions are altered in soybean roots during *M. phaseolina* colonization

Since a large number of CAZymes were detected in our dataset, we investigated the status of soybean root cell walls during *M. phaseolina* infection. The roots of two-week-old soybean plants were infected with the mycelia of *M. phaseolina*, and the cell walls were extracted from the control and infected samples as alcohol-insoluble-residue (AIR). The cell wall samples were further subjected to various biochemical analyses to monitor the changes in the cell walls during infections. First, we characterized the cell walls isolated from the control and infected soybean roots using the Fourier Transform Infrared Spectroscopy (FTIR). This analysis revealed no significant differences in the spectrum profile of control and infected cell wall samples, suggesting no extensive cell wall degradation at the investigated time point (Figure S5). Next, we performed the monosaccharide composition analysis. Here, the neutral sugars from the cell walls were derivatized as alditol acetates (AA) and these residues were analyzed using gas chromatography-mass spectrometry (GC-MS). This analysis has revealed the presence of different neutral sugars, such as arabinose, xylose, mannose, galactose, rhamnose, and glucose, in the cell walls of the control root sample. Of these neutral sugars, the relative abundance of arabinose and xylose was significantly higher than other detected sugars in the control root sample (Figure 2A), indicating that the polysaccharides derived from xylose and arabinose are abundant in the cell walls of soybean roots. Interestingly, the relative abundance of arabinose and xylose was significantly reduced in the infected cell walls compared to the control samples, suggesting the degradation of polysaccharides derived from these neutral sugars (Figure 2B). To gain further insights into structural changes of soybean root cell walls during *M. phaseolina* infections, we performed glycosyl linkages analysis. The neutral sugars from cell walls were derivatized as partially methylated alditol-acetates (PMAA) and were analyzed using GC-MS. Interestingly, this analysis revealed significant differences in the key glycosidic sugar residues in the control and infected samples. The relative abundance of glycosidic sugar residues such as 2-Rha*p*, 5-Ara*f*, 2-Ara*f*, 4-Gal*p*, and 3,6-Gal*p* was significantly reduced in the cell walls of infected root samples compared to control samples (Figure 2C and Table S4). Meanwhile, the relative abundance of 3-Glc*p* was significantly higher in the cell walls of infected samples than in the control. The relative abundance of other detected glycosidic neutral sugar residues was not significantly different between the control and infection samples (Table S4). The glycosidic sugar residues that are detected in lower relative abundance in the infected samples are characteristic linkages derived from cell wall polysaccharides of pectins and hemicellulose. 2-Rha*p* is the main glycosidic sugar present in the main chain of rhamnogalacturonan-1 (RG1) pectins. 5-Ara*f*, 2-Ara*f* and 4-Gal*p* are associated with the arabinogalactans (AGs) linked with RG1 pectins. 3,6-Gal*p* is the characteristic glycosidic linkage present in type II arabinogalactans. Analysis of crystalline cellulose content in the cell wall samples has indicated that the levels of these polymers are reduced in the infected samples compared to the control (Figure S6). These results suggest that the cell wall polysaccharides RG-1, AGs, and cellulose are potentially targeted by CAZymes of *M. phaseolina* during infection.

**Figure 2:**
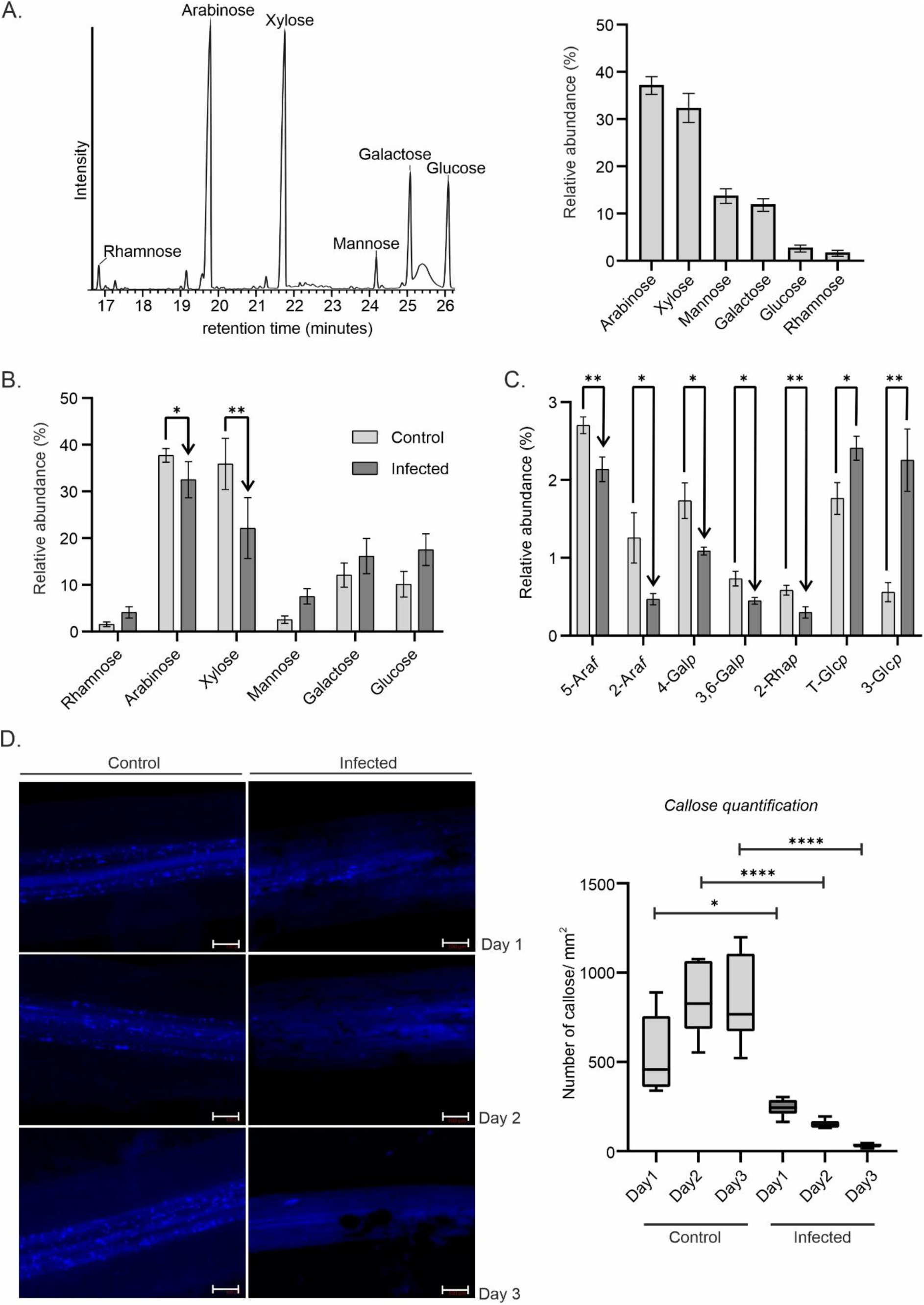
Cell wall composition and callose depositions are altered during *M. phaseolina* infection. (A) Arabinose and xylose are the major neutral sugars detected in the soybean root cell walls. Cell walls were isolated from the soybean roots as alcohol insoluble fraction (AIR) and subjected to monosaccharide composition analysis. The neutral sugars were released from the cell walls by mild TFA (Trifluoroacetic acid) hydrolysis and derivatised as alditol acetate (AA) and the AA residues were analysed using GC-MS. The detected neutral sugars were annotated based on their retention time and the MS spectra of the respective sugar standards. The experiment was conducted with five independent biological replicates. The neutral sugars are expressed as a percentage of the total peak areas of sugars detected. (B) The relative abundance of arabinose and xylose are reduced in the soybean root cell walls infected with *M. phaseolina*. The relative abundance of the neutral sugars detected in the control and infected root AIR samples is expressed as a percentage of the total peak areas of sugars detected. The statistical significance was determined using two-tailed t-test (*: p-value ≤ 0.05, **: 0.001 < p-value ≤ 0.01). The experiment was conducted with five independent biological replicates. (C) The relative abundance of characteristic glycosyl linkages derived from arabinogalactan and the rhamnogalacturonan 1 (RG1) are reduced in the soybean root cell walls infected with *M. phaseolina*. Glycosyl linkage analysis was performed with the AIR samples isolated from control and *M. phaseolina-* infected soybean roots. The sugar residues were derivatized as partially methylated alditol acetates (PMAA) and the residues were analysed using GC-MS. The glycosyl linkages were annotated based on their retention time and the publicly available CCRC spectra library. The glycosyl linkages with significant differences in relative abundance between the control and infected AIR samples are represented. The statistical significance was determined using two-tailed t-test (*: p-value ≤ 0.05, **: 0.001 < p-value ≤ 0.01). (D) The callose levels in the soybean roots are reduced during *M. phaseolina* infection. Aniline blue staining was performed on soybean roots collected at 1 dpi, 2 dpi, and 3 dpi. The stained lateral roots were observed under an epifluorescence microscope using a DAPI filter with an excitation wavelength of 370 nm and an emission wavelength of 509 nm (magnification= 10X, scale =100µm). Callose depositions were quantified using the trainable weka segmentation v3.3.4 plugin of ImageJ 1.54f. The experiment was conducted with six independent biological replicates, two replicates per plant in a completely randomized experimental design. The statistical significance was determined using two-tailed t-test (*: p-value ≤ 0.05, ****: p-value ≤ 0.0001).

3-Glc*p* is the characteristic sugar residue of α-1,3 or β-1,3 glucan polysaccharides. Hence, the higher relative abundance of 3-Glc*p* detected in the infected root samples might be derived from the cell walls of *M. phaseolina* or soybean root callose depositions. Callose deposition is an important immune response displayed by plants to prevent the colonization of fungal pathogens. To investigate if callose depositions are increased in the soybean roots during *M. phaseolina* infections, the control and infected root samples were stained with aniline blue, and the callose was visualized using epifluorescent microscopy. Surprisingly, the number of callose depositions was significantly lower in the infected roots compared to the control at 3 dpi (Figure S7). The detection of a high number of callose depositions in the control roots might be due to the conditions we used for the infection assays. Next, we monitored the dynamics of callose deposition by staining the root samples collected at early time points of infection. Remarkably, the number of callose depositions was gradually reduced during *M. phaseolina* infection, whereas the callose depositions in the control roots remained unchanged at these time points (Figure 2D). These data indicate that callose depositions are degraded during *M. phaseolina* infection. Furthermore, the increase in the relative abundance of 3-Glc*p* in the infected samples is not due to the callose deposition but rather derived from the cell walls of *M. phaseolina*.

### Sequence-unrelated structurally similar (SUSS) effectors and effectors with novel structural folds are secreted by *M. phaseolina* in soybean root apoplast

Plant pathogens secrete protein effectors in apoplast to manipulate the host to favor their colonization process. To investigate the repertoire of candidate effectors secreted by *M. phaseolina*, we analyzed 228 *M. phaseolina* proteins detected in the soybean root apoplast using the EffectorP 3.0 machine learning tool (Sperschneider and Dodds 2022). This analysis has revealed that nearly 23% of proteins (49 proteins) are predicted as candidate apoplastic effector proteins, and one protein was predicted as a cytoplasmic effector (Figure 3A). Pfam analysis has revealed that these 49 candidate apoplastic effectors were diverse and belong to various enzyme classes such as glycosyl hydrolases (eight proteins), polysaccharide lyases (seven proteins), serine hydrolases (three proteins), proteases (three proteins) and other proteins (15 proteins). Interestingly, nine candidate apoplastic effectors could not be classified due to a lack of significant sequence-level similarities, and the conserved domain information for these proteins was not available (Figure 3A). To investigate further if these nine effectors have structural homology with other fungal effector proteins, we performed the AlphaFold (AF) analysis. The candidate proteins were modelled as monomers using the AF2 program, and template-free AF2 modelling was implemented in the local colabfold pipeline with 5X models for each unique pair (Figure 3B). The resulting AF models were relaxed using an amber force field for energy minimization, and the relaxed structural models were further analyzed (Figure 3B). The top-ranked structural model for each candidate effector was compared with the experimentally resolved protein crystal structures in the PDB100 database using the Foldseek search program. Since *M. phaseolina* belongs to the division Ascomycota of the fungal kingdom, we restricted our Foldseek search to this division. Interestingly, this analysis has revealed that seven proteins shared structural homology with the proteins from the members of the Ascomycota division with a significant TM score > 0.5 and RMSD values ≤ 4.0 (Table S5) and two proteins (UniProt IDs: K2R751 and K2RET3) that did not share similarities at structural levels to proteins from Ascomycota division (Figure S8 A,B). Interestingly, when searching the structural model of these proteins against the AlphaFold protein structure database (AlphaFold DB) revealed that these proteins have structural similarities with proteins from the Ascomycota fungal division, suggesting that these are potentially uncharacterized family of SUSS effectors and their structural folds are evolutionarily expanded across the members of this division (Table S6). Notably, these proteins did not share any structural similarities with any of the structurally resolved proteins of other fungal divisions, indicating that the structural folds in these proteins are possibly novel. Of the seven proteins with predicted structural homologs, five proteins (Uniprot IDs: K2RZ98, K2RDS7 K2QS10, K2RZH1, and K2R1W5) shared similarities with effectors that are secreted by plant-associated fungi and two proteins (Uniprot IDs: K2RGG5 and K2RF41) showed significant structural similarities with proteins from *Neurospora crassa* and *Saccharomyces cerevisiae*, respectively (Figure S9). The protein K2RZ98 shared structural similarity to the Tsp1 effector of *Trichoderma virens* Gv29-8, a beneficial plant fungus with a significant TM-score of 0.801 and an RMSD of 1.23 Å (Figure 3C). K2RZ98 displayed a distinct β-barrel fold unique to the Tsp1 effector (Gupta et al. 2021) (Figure 3C). The protein K2RDS7 shared structural similarity to Zt-KP6-1 effector from *Zymoseptoria tritici*, a plant fungal pathogen with a TM-score and RMSD score of 0.503 and 2.89 Å, respectively. The K2RDS7 has a characteristic KP6-like fold consisting of two signature α-helices and a variable number of β-sheets (3-5) (Rocafort et al. 2022). Remarkably, three candidate effector proteins, K2RZH1, K2R1W5 and K2QS10, shared structural similarity to the MoErs1 protein with a TM score > 0.5 and RMSD scores in the range of 3-3.8 Å. MoErs1 is an effector secreted by *Magnaporthe oryzae*, a fungal pathogen that causes the rice blast disease. These three proteins were predicted to significantly share the characteristic β-trefoil fold displayed by the MoErs1 effector (Y. Liu et al. 2024). Taken together, our AF2 analysis has revealed that SUSS effectors and effectors with novel structural folds are secreted by *M. phaseolina* in soybean root apoplast during infections.

**Figure 3:**
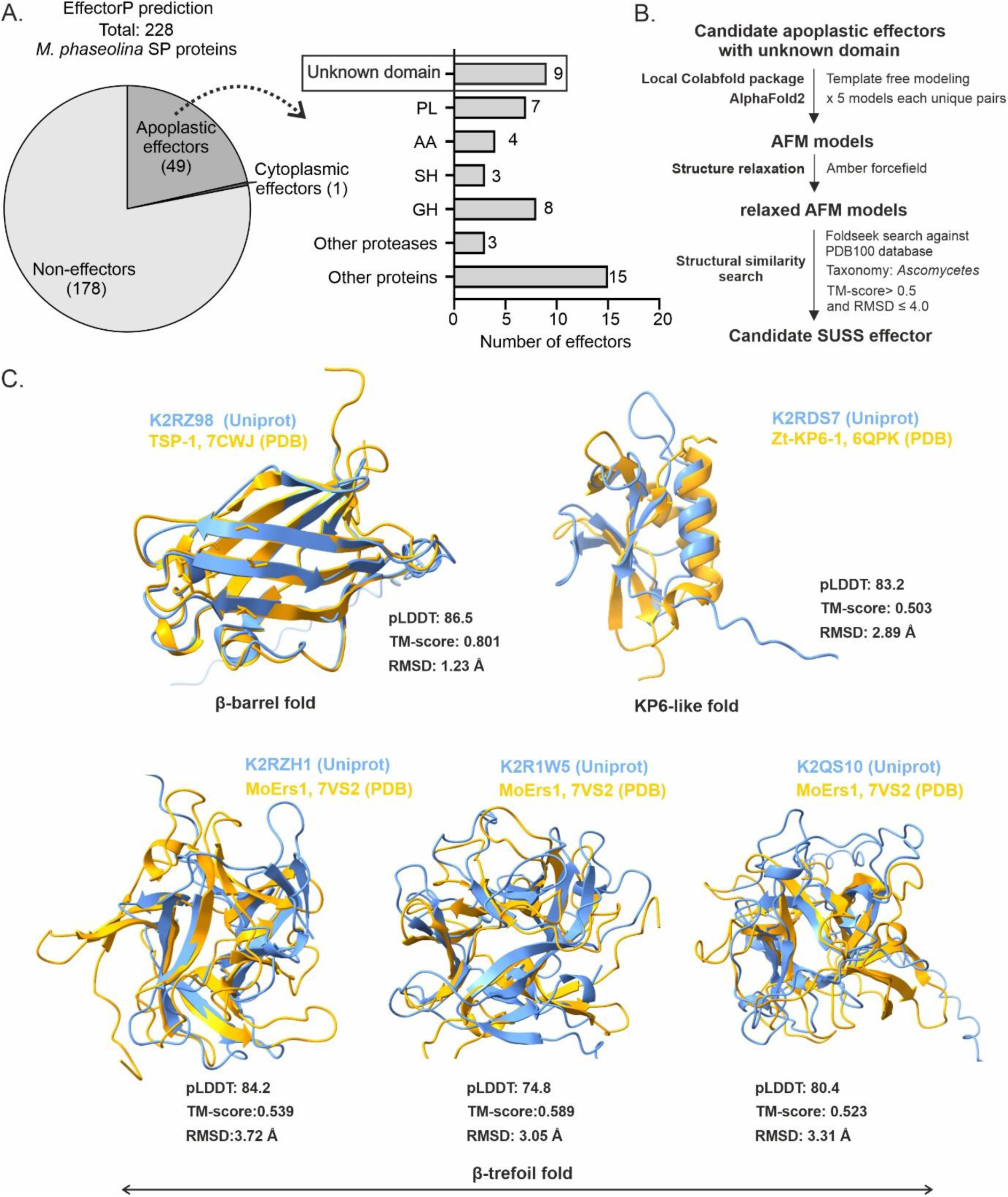
*M. phaseolina* secretes SUSS effectors and effectors with novel structural folds in the soybean root apoplast during infection. (A) Apoplastic effectors are secreted by *M. phaseolina*. *M. phaseolina* proteins carrying signal peptide were analysed using EffectorP Fungi 3.0 (https://effectorp.csiro.au/) for effector prediction. The predicted apoplastic effector candidates were classified based on their domain architecture using Pfam analysis. (B) Pipeline for predicting SUSS effector candidates using the AF2 and Foldseek program. The candidate apoplastic effectors with unknown domain were modelled using AF2 implemented in colabfold pipeline without template. The AF2 modelled effector candidates were screened against PDB100 database using Foldseek search with taxonomy restriction to Ascomycetes. The protein candidates with TM-score >0.5 were considered structurally similar. (C) AF2 predicts SUSS effector candidates secreted by *M. phaseolina*. Superposition (structural alignment) of AF2 predicted putative SUSS effectors from *M. phaseolina* to experimentally resolved 3D tertiary structures available in PDB100 database were performed using Foldseek search and matchmaker module of chimera molecular viewer. The predicted putative SUSS effectors are represented in blue whereas crystal structures of known effectors are represented in yellow. The protein K2RZ98 shows similarity to Tsp-1 effector from a beneficial fungi *Trichoderma virens* (PDB ID: 7CWJ). K2RDS7 protein shows folding similar to Zt-KP6-1 effector (PDB ID: 6QPK) from *Zymoseptoria tritici*. The proteins K2RZH1, K2R1W5, and K2QS10 show similarity to MoERS-1 effector from *Magnaporthe oryzae* (PDB ID: 7VS2). SP, signal peptide, PL, polysaccharide lyases, AA, auxiliary activities, SH, serine hydrolases, GH, glycosyl hydrolases, AF2, AlphaFold 2, SUSS, sequence unrelated structurally similar, PDB, Protein data bank, pLDTT, predicted local distance difference test, TM-score, template modelling scores, RMSD, root mean square deviation.

### AFM analysis combined with molecular dynamics (MD) reveals the interaction of MoErs1-like effector (K2QS10) with secreted cysteine protease (I1JTM0)

MoErs1 is reported to be a species-specific effector secreted by *M. oryzae* to inhibit the activities of OsRD21, a rice cysteine protease (Y. Liu et al. 2024). However, our AF2 analysis revealed that the three effectors from *M. phaseolina* shared structural homology with MoErs1, indicating that the MoErs1-like effectors are present in *M. phaseolina*. This prompted us to investigate these proteins further. To investigate if the MoErs1-like secreted effectors from *M. phaseolina* can target soybean cysteine proteases, we performed *in silico* pulldown with the AlphaFold Multimer (AFM) platform. The pipeline for our analysis involves modelling ligand and bait proteins as heterodimers using MMseqs2 for the template search (Figure 4A) (Mirdita et al. 2019). The AFM-modelled structures were relaxed using an amber force field, and the top five models were considered (Mirdita et al. 2022). To investigate if AFM can distinguish between existing and non-existing protein complexes, first, we modelled a well-characterized and experimentally validated protease-inhibitor complex of P69B, a serine protease from tomato against Epi1A, a serine protease inhibitor from *P. infestans* (Tian et al. 2005). Our AFM pipeline with MMseqs2 predicted the P69B-Epi1A complex at high quality with pLDDT score of 82.5, pTM score of 0.77, ipTM score of 0.78 and confidence score of 79% (Figure S10). Interestingly, replacing Epi1A with EpiC2B, a cysteine protease inhibitor from *P. infestans* (Tian et al. 2007), our AFM pipeline predicted the P69B-EpiC2B complex at high quality with a pLDDT score of 83.6, pTM score of 0.79 and notably a low significant ipTM score of 0.26. This result indicates the ability of our AFM pipeline to distinguish between existing and non-existing protein complexes (Figure S10). These results are similar to the AFM prediction performed recently for these complexes (Homma et al. 2023; Mooney and van der Hoorn 2024). Next, we followed the above AFM pipeline for screening MoErs1-like secreted effectors against soybean cysteine proteases with a stringent scoring threshold (pTM+ipTM > 1.1, pLDDT > 85, and model confidence score > 75%) based on the P69B-Epi1A complex AFM scores and previously reported scoring schemes for confident AFM predictions (Tian et al. 2005) (Figure 4A). We used three MoErs1-like secreted effectors (Uniprot IDs: K2RZH1, K2R1W5, and K2QS10) as ligands against five soybean cysteine proteases (Uniprot IDs: I1MXL3, I1MHG0, I1MER7, I1JTM0 and I1M2Y6) as baits. These cysteine proteases were detected in the soybean root apoplast both in control and infected samples (Table S7). In parallel, we performed the AFM analysis of MoErs1 (ligand) against the OsRD21 cysteine protease (bait). The pro-inhibitory domain present in the N-terminus of cysteine proteases was removed before analysis as recommended for AFM analysis of cysteine proteases (Homma et al. 2024). The resultant structural models predicted by AFM were screened using ipTM+pTM score, pLDDTscore, and model confidence to select the top interacting partners (Figure 4A). Interestingly, our AFM analysis has revealed several interaction models for MoErs1-like effectors with soybean cysteine proteases with significant ipTM+pTM and pLDDT scores (Figure 4B). Next, we employed model confidence score selection with > 75% cut-off to select the top interaction models (Figure 4C). Remarkably, of the three MoErs1-like effectors, one MoErs1-like effector (UniprotID. K2QS10) displayed significant interaction with the secreted soybean cysteine protease (UniprotID. I1JTM0) (Figure 4B,C). The interaction model predicted by AFM has a significant ipTM+pTM score of 1.60, a high pLDDT score of 84.36, and a model confidence score of 81% (Figure 4D). These cut-off values are comparable with the AFM predictions for MoErs1 and OsRD21 with an ipTM+pTM score of 1.65, supported by a high pLDDT score of 89.8 and a confidence score of 85.8% (Figure 4D). Notably, the loop regions of the K2QS10 displayed significant interaction with the amino acids present in the active site of I1JTM0 and these interactions are comparable to AFM models of MoErs1-OsRD21 complex (Figure 4E). In general, the N-terminus of the plant cysteine proteases carries a self-inhibitory regulatory pro-domain that blocks the active site of these enzymes (Hou et al. 2018). Remarkably, performing the AFM analysis with the pro-inhibitory domain included in the cysteine protease resulted in non-significant models with low ipTM+pTM and pLDDTscore and low model confidence scores (Figure S11A,B). Therefore, this result suggests that the MoErs1-like effector, K2QS10, secreted by *M. phaseolina*, might serve as an active site-directed cysteine protease inhibitor.

**Figure 4:**
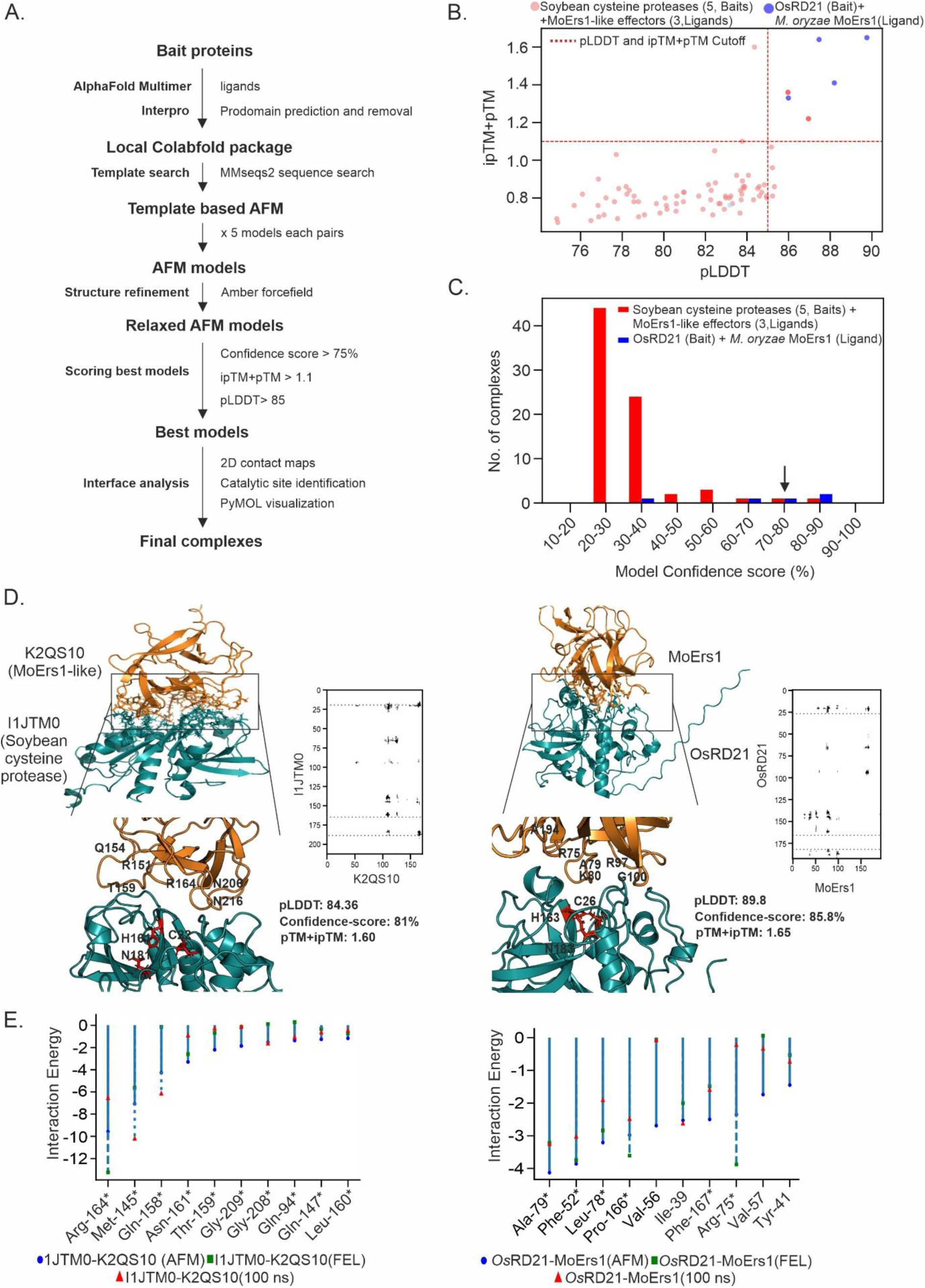
AFM analysis predicts significant interaction of a MoErs1-like effector (UniprotID. K2QS10) with a soybean cysteine protease (UniprotID. IJTM0). (A) The workflow for predicting interactions between proteins (bait and ligands) using AFM. The pipeline involves modelling ligand and bait proteins as heterodimers using the template from MMseqs2 search. The AFM-modelled structures were relaxed using an amber force field, and the top five models were considered. The resultant structural models predicted by AFM were screened using the stringent ipTM+pTM score (> 1.1), pLDDT score (> 85), and model confidence (> 75%) to select the top interacting partners. The protein complexes satisfying this threshold were further analyzed for interfacial contacts using 2D contact maps, and the interactions were visualized using the PyMOL software. (B) AFM analysis reveals interactions of MoErs1-like effectors with soybean cysteine proteases. The cysteine proteases detected in the soybean root apoplast and the rice RD21 protease (OsRD21) were screened against MoErs1-like effectors secreted by *M. phaseolina* and the MoErs1 effector from *M. oryzae* using AFM. This analysis was implemented in the local colabfold pipeline using the ipTM + pTM and pLDDT scores. 120 AFM models are distributed over the full pTM+ ipTM score and pLDDT range, with a subpopulation of high confidence AFM model predictions with pTM+ ipTM score > 1.1 and pLDDT > 85 (top right corner). The top-ranked AFM models are colored in darker shades of red and blue (Red represents the models of MoErs1-like effector and soybean cysteine proteases, and blue represents the models of MoErs1 and OsRD21). (C) AFM predicts the interaction of MoErs1-like effector (UniprotID. K2QS10) with a soybean cysteine protease (UniprotID. IJTM0) with a high model confidence score. The AFM interaction model with a confidence score > 75% is indicated with an arrow. (D) The MoErs1-like effector (UniprotID. K2QS10) and MoErs1 interact in close proximity to the active sites of respective cysteine proteases. The AFM-predicted interaction models were visualized using the PyMOL software and presented as a cartoon backbone 3D representation. The heterodimer interface of interaction between K2QS10 (MoErs1-like effector) and I1JTM0 (soybean cysteine proteases) and MoErs1 and OsRD21 is zoomed and represented as an insert. The putative active site catalytic triads in cysteine proteases were predicted using InterPro and represented as red sticks. The amino acid residues forming hydrogen bonds near the active site are represented as single-letter codes with residue numbers. The interfacial contacts in the protein complexes are presented as 2D contact maps. (E) The loop region of MoErs1-like effector and MoErs1 displays energetically favourable interactions with the respective cysteine proteases. The per-residue decomposition plots of MoErs1-like effector and MoErs1 with the respective cysteine protease complex were using the HawkDock tool. The decomposition plots represent the interaction energy of amino acid residue in the loop region of MoErs1-like effector and MoErs1. The dark blue dot represents the protein complex from the AFM model, the green square represents the low energy conformer obtained through FEL analysis, and the red triangle represents the conformer from 100^th^ ns in the MD trajectories. The light blue line in the plot indicates the range of interaction energy values for each amino acid, and the blue line is dotted in the plot for easy visualization of other data points. The symbol * represents amino acids at loop regions of MoErs1 and MoErs1-like effectors that interact with the respective cysteine proteases. pLDTT, predicted local distance difference test, pTM, predicted template modelling, ipTM, interface predicted template modelling, AF2, AlphaFold 2, SUSS, Sequence unrelated structurally similar, PDB, Protein data bank, TM-score, template modelling scores, 2D, two-dimensional, FEL, Free energy landscape, AFM, AlphaFold multimer, ns, nano seconds.

To further validate the predicted AFM interactions, we performed MD simulations of the MoErs1-like effector and soybean cysteine protease complex. In parallel, we performed these analyses with MoErs1 and OsRD21 cysteine protease complex. All-atom MD simulations were performed using the OPLS-AA force field for a 100 nanoseconds (ns) time scale. Root mean square deviation (RMSD) analysis was performed to evaluate the reliability and stability of the protein complexes. Additionally, the trajectories obtained from the MD simulations were subjected to MM-PB/GBSA calculations to evaluate the binding affinities of the complexes. The RMSD analysis of the MD trajectories indicated that the K2QS10 and the cysteine I1JTM0 complex exhibited a constant RMSD after 40 ns, indicating that the protein-inhibitor complex reached equilibrium during the MD run (Figure S12A). The distance between these protein-inhibitor complexes remained constant after 80^th^ ns, indicating that there is no conformational shift at the contact interface (Figure S12B). This is supported by the number of hydrogen bonds calculated at the interface and 2D contact maps showing contact frequency during 80-100th ns (Figures S12C,D). The MM-PB/GBSA calculations for binding affinity have indicated that the protein complex is energetically favourable with negative ΔG values (Table S8). Next, we performed a per-residue decomposition analysis using the HawkDock tool to identify the significant interacting amino acid residue at the contact interface. For this analysis, first, we performed FEL analysis on the MD trajectories to obtain low-energy conformation of K2QS10 and I1JTM0 complex (Figures S12E). Next, we integrated the top interaction model predicted by AFM, the low energy conformer obtained through FEL analysis, and the conformer from 100^th^ ns in the MD trajectories for per-residue decomposition analysis. This analysis has revealed significant interactions of amino acid residues present in the loop regions of the K2QS10 with I1JTM0 (Figure 4E). The amino acid residues present in the loop region of the K2QS10 displayed similar interaction energy values with I1JTM0 in all three structural models used for the analysis. This result suggests that the loop regions in the K2QS10 might have a crucial role in inhibiting I1JTM0. This is consistent with the crystal structures of cysteine protease inhibitors such as chagasin and cystatin, where the loop region interacts with the active site to inhibit the activities of cysteine proteases (Bode et al. 1988; Redzynia et al. 2009). Notably, the MD simulations and binding calculation analyses performed for the K2QS10-I1JTM0 complex were comparable with the MoErs-1-OsRD21 complex (Figure 4E and Figure S12A-E). Taken together, the AFM analysis together with the MD simulation indicates that the secreted MoErs1-like effector, K2QS10, might have a role as an apoplastic cysteine protease inhibitor.

### Quantitative proteome analysis reveals the dynamics of soybean apoplastic proteins during *M. phaseolina* infection

In our apoplastic proteomic dataset, we detected 255 secreted proteins from soybean that carried a signal peptide. Pfam analysis of 255 secreted soybean proteins has revealed that the majority of these proteins were CAZymes (98 proteins, 38%) followed by proteases (27 proteins, 12%) and protease inhibitors (16 proteins, 7%) (Figure 5A and Table S7). The other 114 proteins were diverse and included CAP proteins (Cysteine-rich secretory protein [CRISP], Antigen 5 [Ag5], pathogenesis-related 1 protein, three proteins), fasciclin-like arabinogalactans (FLAs) (nine proteins), dirigent proteins (seven proteins), thioredoxin (four proteins) and other proteins (Table S7). Of the 255 secreted proteins, 241 proteins were detected in both control and infected samples (Table S9). Eight proteins were detected as uniquely in control samples, and six proteins were detected uniquely in infected samples (Figure S13). To monitor the dynamics of soybean apoplastic proteins during *M. phaseolina* infection, we performed a volcano plot and z-score analysis. In the volcano plot analysis, -log_10_ p-values of the 241 proteins detected in both control and infected samples were plotted against their respective log_2_ fold change of differential abundance values (Figure 5B). Z-scores of the 241 proteins were calculated by normalizing the LFQ values with their standard deviation, and the z-score values were represented as a heat map to visualize the differentially regulated proteins (Figure S14). These analyses have revealed that 95 soybean apoplastic proteins were differentially regulated during *M. phaseolina* infection. 76 proteins were upregulated, 19 proteins were downregulated, and 146 proteins remained unregulated (Figure 5B). Gene ontology analysis of the 95 differentially regulated proteins has revealed that the proteins that are associated with several molecular functions and biological processes are detected in our dataset. The proteins associated with lactoperoxidase activity are highly enriched in the upregulated proteins compared to the downregulated proteins (Figure 5C). Furthermore, the proteins associated with several molecular functions such as peptidase inhibitor, heme binding, chitinase activity, alpha-mannosidase activity, chitin-binding, serine peptidase activity, and Flavin Adenine Dinucleotide (FAD) binding are uniquely detected in the upregulated proteins (Figure 5C). Interestingly, the proteins that are associated with the fatty acid binding are uniquely detected only in the downregulated proteins. In addition, the proteins associated with biological processes such as hydrogen peroxide catabolic process and cellular oxidant detoxification are highly enriched in the upregulated proteins compared to the downregulated proteins (Figure 5C). The proteins associated with several biological processes, such as the mannose metabolic process, amino sugar catabolic process, phenylpropanoid biosynthetic process, and cell wall catabolic process, are uniquely detected in the upregulated proteins (Figure 5C). Pfam analysis of 76 upregulated secreted proteins of soybean has revealed these proteins are classified into CAZymes (27, 36%), proteases (seven, 9%), protease inhibitors (nine, 11%), and other diverse proteins (33, 43%) (Figure 5D). The detected CAZymes belonged to various enzyme classes, such as glycosyl hydrolases (GH), carbohydrate esterases (CE), auxiliary activity (AA) and carbohydrate-binding module (CBM). Among the seven upregulated proteases, six proteins were serine proteases from families S8, S10, and S28, and one cysteine protease from family C26. Interestingly, three only leucine-rich repeat-containing proteins (LRR) and two expansin proteins were upregulated during *M. phaseolina* infection. These proteins are previously reported to act as susceptibility factors in other plant species during fungal infections (Abuqamar et al. 2013; Chen et al. 2021). Notably, all the upregulated protease inhibitors (Nine proteins) belonged to Kunitz-type serine protease inhibitors. Pfam analysis of 19 downregulated proteins of soybean has revealed these proteins are classified into nine CAZymes from different families, four proteases, and six diverse proteins that include trichome birefringence-like (TBL) protein (one protein), fasciclin-like arabinogalactan-protein (FLA) (two proteins), lipid transfer proteins (two proteins), and lipid recognition protein (one protein) (Figure 5D). The categorization of the differentially regulated proteins based on the pathogenesis-related proteins (PR) has revealed PR proteins belonging to families PR-2, PR-3, PR-5, and PR-9 are upregulated, and the PR proteins belonging to families PR2, PR9, and PR14 are downregulated during *M. phaseolina* infections (Figure 5D). Taken together, this quantitative proteome analysis of the soybean-secreted proteins revealed differential regulation of several apoplastic proteins during *M. phaseolina* infection.

**Figure 5:**
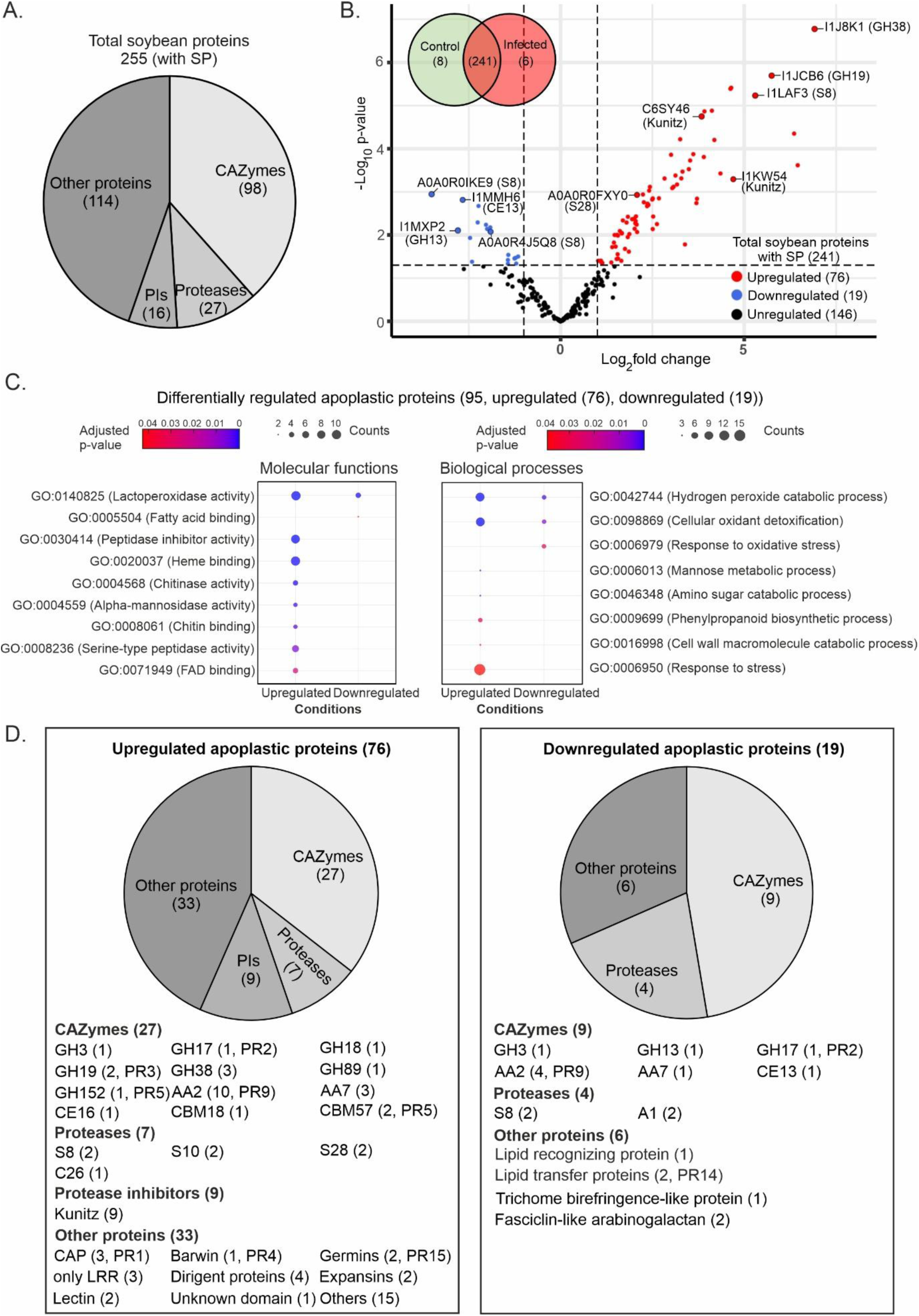
Quantitative proteome analysis reveals the dynamics of soybean-secreted proteins in root apoplast during *M. phaseolina* infection. (A) A diverse class of soybean-secreted proteins are detected in the root apoplast. 255 soybean proteins with signal peptides were classified based on their domain architecture using dbCAN3 and Pfam analysis. (B) Differential regulation of soybean apoplastic proteins upon *M. phaseolina* infection. Soybean apoplastic proteins detected in control and infected apoplastic fluids are represented as a Venn diagram. Soybean apoplastic proteins are ranked in a volcano plot according to their statistical P value (y-axis) as -log 10 and their relative Label-Free Quantification (LFQ) values (Log_2_ fold change, x-axis) between control *and M. phaseolina-*infected soybean root samples (n=3) using the EnhancedVolcano package in the R program. Log p-value signifies its statistical significance, i.e., the lower the p-value, the greater the significance of the value. Log_2_ fold change was calculated to understand the differential regulation of the apoplastic proteins in the infected apoplast compared to the control apoplast. The upregulated proteins with log_2_ fold change > 1 and p-value < 0.05 are represented in red colour, downregulated proteins with log_2_ fold change < -1 and p-value < 0.05 are represented in blue colour and the rest of the proteins are represented as black dots. The proteins with bigger dots represent the top two highly regulated proteins of the major classes. (C) Gene ontology analysis of the differentially regulated soybean apoplastic proteins. GO annotation was performed for upregulated and downregulated soybean apoplastic proteins using g:GOSt module of g:Profiler based on molecular functions and biological processes. Statistical significance was calculated using the Fisher one-tailed test in combination with the Benjamini-Hochberg with an FDR threshold of ≤0.05 to correct for multiple testing. The colour code signifies the adjusted p-value and the bubble size denotes the counts of genes. The adjusted p-value is a statistical measure denoting the significance of enrichment, where a low value indicates the result is less likely to occur by chance. GO IDs represent taxonomy that statistically determines the involvement of those sets of genes in the specific biological processes and molecular functions as indexed in QuickGo (https://www.ebi.ac.uk/QuickGO/). (D) Classification of differentially regulated soybean apoplastic proteins. Significantly upregulated and downregulated soybean proteins were classified based on their domain architecture using Pfam analysis. The proteins of their respective classes are listed. SP, signal peptide, GH, glycosyl hydrolases, PIs, protease inhibitors, CE, carbohydrate esterases, CBM, carbohydrate binding module, AA, auxiliary activities, CAP, Cysteine-rich, Antigen-5, PR-1, LRR, Leucine-rich repeats, S#, serine proteases, C#, cysteine proteases, A#, aspartic proteases, PR proteins, pathogenesis-related proteins. #, number

### AFM analysis combined with activity-based protein profiling (ABPP) reveals interactions of soybean Kunitz with serine proteases

Plant protease inhibitors are involved in the regulation of immune protease activities. In our proteome dataset, we have detected 16 protease inhibitors in the soybean root apoplast. Of these 16 protease inhibitors, 15 belong to Kunitz-type protease inhibitors (Kunitz proteins), and one belong to cysteine protease inhibitor. Interestingly, among these detected protease inhibitors, only the Kunitz proteins were upregulated in abundance during *M. phaseolina* infections (Figure 6A). Nine Kunitz proteins (Uniprot IDs: I1LVB0, C6T2D3, I1MI59, C6T586, I1KYW5, I1KYX0, C6SY46, I1KW54, and I1LVB1) were significantly abundant in the infected samples compared to the control and one Kunitz protein (Uniprot ID: I1KYX1) was uniquely detected only in the infected samples (Figure 6A and Figure S13). To investigate the structural relationship between the upregulated Kunitz proteins, we modelled the ten Kunitz proteins with AF2 as monomers and assessed their structural similarity using the DALI web server (Holm 2022). This analysis has revealed that the detected soybean Kunitz can be classified into two groups: Group 1 consists of six Kunitz proteins (Uniprot IDs: I1MI59, I1KW54, I1LVB1, I1LVB0, C6SY46, C6T2D3) and group 2 consists of 4 Kunitz proteins (Uniprot IDs: I1KYW5, C6T586, I1KYX0, I1KYX1) (Figure S15). This result indicates that Kunitz proteins detected in soybean root apoplast are structurally diverse. Plant Kunitz-type protease inhibitors have been predominately investigated for their role in regulating the activities of serine proteases derived from insects and humans (Mehmood et al. 2024; Okedigba et al. 2023). To investigate the potential targets of the Kunitz proteins that are upregulated in our dataset, we employed AFM analysis to predict interactions of these Kunitz proteins with randomly selected secreted serine proteases detected in soybean root apoplast. We followed the pipeline and the stringent cut-off scores that we used for the AFM analysis of soybean cysteine proteases and MoErs1 (Figure 6B). In the AFM analysis, the ten soybean Kunitz proteins (Nine upregulated and one uniquely expressed) were used as ligands against ten soybean serine proteases and seven *M. phaseolina* serine proteases that were randomly selected from our dataset (Tables S10 and S11). The soybean serine proteases belonged to families S8, S10, and S28, whereas the *M. phaseolina* serine proteases belonged to families S8, S9, and S10 (Tables S10 and S11). Interestingly, our AFM analysis of the upregulated Kunitz proteins against the secreted serine proteases has revealed the interaction of Kunitz proteins with serine proteases of soybean and *M. phaseolina* with significant ipTM+pTM, pLDDT, and high model confidence scores (Figure 6 C, D). Notably, the AFM analysis of soybean Kunitz proteins with randomly selected secreted chitinases detected in our dataset revealed no significant interaction models for the complexes (Figure S16). This result indicates the reliability of our AFM prediction of soybean Kunitz with the serine proteases. The Kunitz proteins from group 1 and group 2, such as I1KW54, I1MI59, I1LVB1, L1LVB0, C6T2D3, and C6T586, are predicted to interact with serine proteases of both soybean and *M. phaseolina* (Tables S10 and S11). The group 2 Kunitz protein, I1KYW5, is predicted to interact only with a serine protease of *M. phaseolina* but not soybean serine proteases. Meanwhile, the other Kunitz protein from group 2, I1KYX1, interacts only with a serine protease of soybean but not *M. phaseolina* serine proteases. The group 1 Kunitz protein, C6SY46, and group 2 Kunitz protein, I1KYX0, did not interact with any of the analyzed serine proteases from soybean or *M. phaseolina* (Tables S10 and S11). Taken together, our AFM analysis indicates distinct interactions of soybean Kunitz proteins with serine proteases from soybean and *M. phaseolina*.

**Figure 6:**
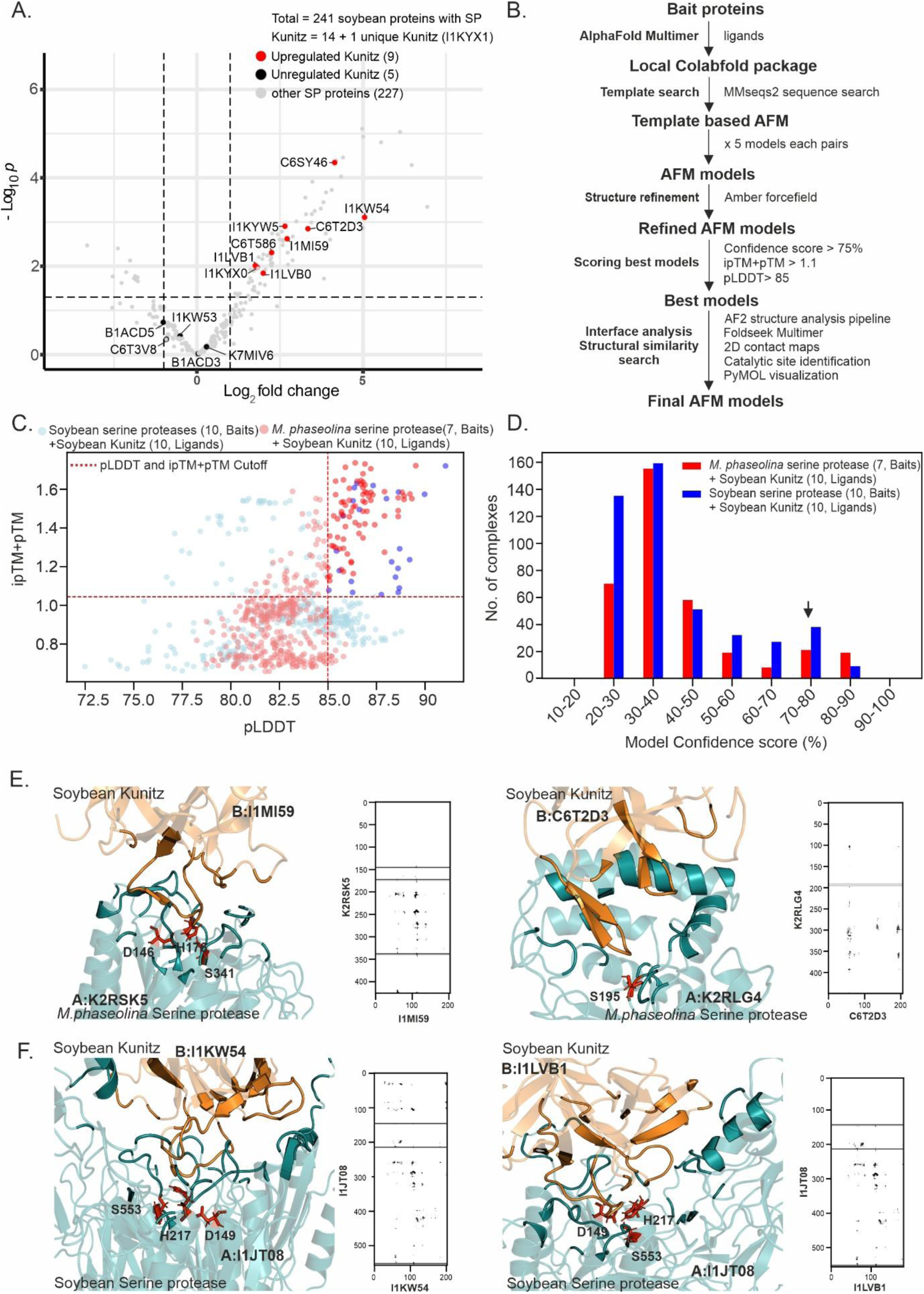
AlphaFold Multimer (AFM) analysis predicts the interaction of soybean Kunitz and serine proteases. (A) Soybean Kunitz proteins are upregulated in soybean root apoplast during *M. phaseolina* infection. Soybean apoplastic proteins are ranked in a volcano plot according to their statistical P value (y-axis) as -log 10 and their relative Label-Free Quantification (LFQ) values (log_2_ fold change, x-axis) between control *and M. phaseolina-*infected soybean root samples (n=3) using EnhancedVolcano package of R software. Log p-value signifies its statistical significance, i.e., the lower the p-value, the greater the significance of the value. Log_2_ fold change was calculated to understand the differential regulation of the apoplastic proteins in the infected apoplast compared to the control apoplast. Red and black dots represent the upregulated and unregulated soybean Kunitz, respectively, and the grey dot represents the other signal peptide containing proteins identified in the soybean apoplast. (B) The workflow for predicting interactions between proteins (bait and ligands) using AFM. The pipeline involves modelling ligand and bait proteins as heterodimers using the template from MMseqs2 search. The AFM-modelled structures were relaxed using an amber force field, and the top five models were considered. The resultant structural models predicted by AFM were screened using the stringent ipTM+pTM score (> 1.1), pLDDT score (> 85), and model confidence (> 75%) to select the top interacting partners. The protein complexes satisfying this threshold were further analyzed for interfacial contacts using 2D contact maps, and the interactions were visualized using the PyMOL software. (C) AFM analysis reveals interactions of soybean Kunitz with serine proteases of soybean and *M. phaseolina*. The upregulated soybean Kunitz in our dataset was screened against the randomly selected ten secreted serine proteases of soybean and seven secreted serine proteases *M. phaseolina* using AFM. This analysis was implemented in the local colabfold pipeline using the ipTM + pTM and pLDDT scores. 120 AFM models are distributed over the full pTM+ ipTM score and pLDDT range, with a subpopulation of high confidence AFM model predictions with pTM+ ipTM score > 1.1 and pLDDT > 85 (top right corner). The top-ranked AFM models are colored in darker shades of red and blue (Red represents the models of *M. phaseolina* serine proteases and Soybean Kunitz, and blue represents the models of Soybean serine proteases and Soybean Kunitz). (D) AFM predicts the interaction of soybean Kunitz with serine proteases of soybean and *M. phaseolina* with a high model confidence score. The AFM interaction models with a confidence score > 75% are indicated with an arrow. (E) The Soybean Kunitz proteins (UniprotIDs: I1MI59 and C6T2D3) interact with the active site of the *M. phaseolina* serine proteases (UniprotIDs: K2RSK5 and K2RLG4). (F) The Soybean Kunitz proteins (UniprotIDs: I1KW54 and I1LVB1) interact with the active site of the soybean serine protease (UniprotID. I1JT08). The AFM-predicted interaction models in Figure 6E and Figure 6F were visualized using the PyMOL software and presented as a cartoon backbone 3D representation. The heterodimer interface of interaction between soybean Kunitz and serine proteases is zoomed and represented. The annotated active site catalytic triads in serine proteases are represented as red sticks. The interfacial contacts in the protein complexes are presented as 2D contact maps. SP, signal peptide, AFM, AlphaFold Multimer, AF2, AlphaFold2, 2D, two-dimensional, pTM, predicted template modelling scores, ipTM, interface predicted template modelling score, pLDDT, predicted local distance difference test score.

Next, we analyzed the AFM-predicted complexes where the active sites and the catalytic residues for serine proteases are well-annotated. Of the several serine proteases that were used in the AFM analysis, the active site catalytic residues for a soybean serine protease (I1JT08) and two fungal serine proteases (K2RSK5 and K2RLG4) are well-annotated in InterPro database. Analysis of these complexes has revealed that these soybean Kunitz proteins interact with the catalytic active site residues of these serine proteases. The Kunitz proteins are reported to inhibit serine proteases through the stearic blockade by binding to the targets with characteristic loop regions (Guerra et al. 2023). A similar loop region could be visualized for the soybean Kunitz proteins in close proximity near the active site of these serine proteases, suggesting the Kunitz proteins might act as an active site-directed inhibitor of serine protease from soybean and *M. phaseolina* (Figure 6E, F). These predicted AFM models of the analyzed complexes are similar to the recent crystal structures of soybean Kunitz and insect serine protease complexes (Mehmood et al. 2024). To further validate these AFM predictions, we performed competitive activity-based protein profiling (ABPP) with the apoplastic fluids. The apoplastic fluids were collected from the control and *M. phaseolina-*infected soybean roots at 3 dpi. The apoplastic fluids were preincubated with different concentrations of the commercially available Kunitz inhibitor of soybean origin. The preincubated proteomes were treated with the fluorophosphate-alkyne probe (FP-alkyne). FP-alkyne is an active site-directed probe that targets several serine hydrolases, including serine proteases (Kaschani et al. 2009). The FP-alkyne-labeled proteomes were subsequently subjected to click chemistry to tag the rhodamine fluorescent moiety. The fluorescently labelled proteins were separated by SDS-PAGE and detected by in-gel fluorescence. This analysis revealed several active serine hydrolases at various molecular weights were detected in the infected and control apoplastic fluid samples compared to the no probe control (Figure 7A,B). Interestingly, the intensities of three bands (band 2, band 3, and band 4) at approx. 75 kDa, 70 kDa, and 55 kDa, respectively, were reduced in the infected samples preincubated with the soybean Kunitz (Figure 7A). Furthermore, the intensities of three bands (band 2, band 4, and band 5) at approx. 75 kDa, 55 kDa, and 50 kDa, respectively, were reduced in the control samples preincubated with the soybean Kunitz (Figure 7B). These results suggest the soybean Kunitz protein can interact with the active site of serine proteases in the control and infected apoplastic fluids. To investigate if the soybean Kunitz can inhibit the serine proteases from *M. phaseolina*, we performed competitive ABPP using the soybean Kunitz and the culture filtrate of *M. phaseolina.* This analysis has revealed that intensities of two bands (band 1 and band 2) at approx. 75 kDa were reduced in *M. phaseolina* culture filtrate samples preincubated with soybean Kunitz. This result suggests that soybean Kunitz protein can also interact with the active site of serine proteases from *M. phaseolina*. Interestingly, we also detected increased labeling intensities of bands in the apoplastic fluid samples and *M. phaseolina* culture filtrates preincubated with soybean Kunitz (Figure 7A-C). The increased intensities might be due to the putative roles of Kunitz in activating serine hydrolases, or indirect activation of some serine hydrolases due to the inhibition of serine protease by Kunitz in the labeling reaction. The exact reason for increased labeling upon Kunitz’s pre-treatment requires further investigation. Taken together, our AFM analysis combined with ABPP reveals the interaction of soybean Kunitz with serine proteases of soybean and *M. phaseolina*.

**Figure 7:**
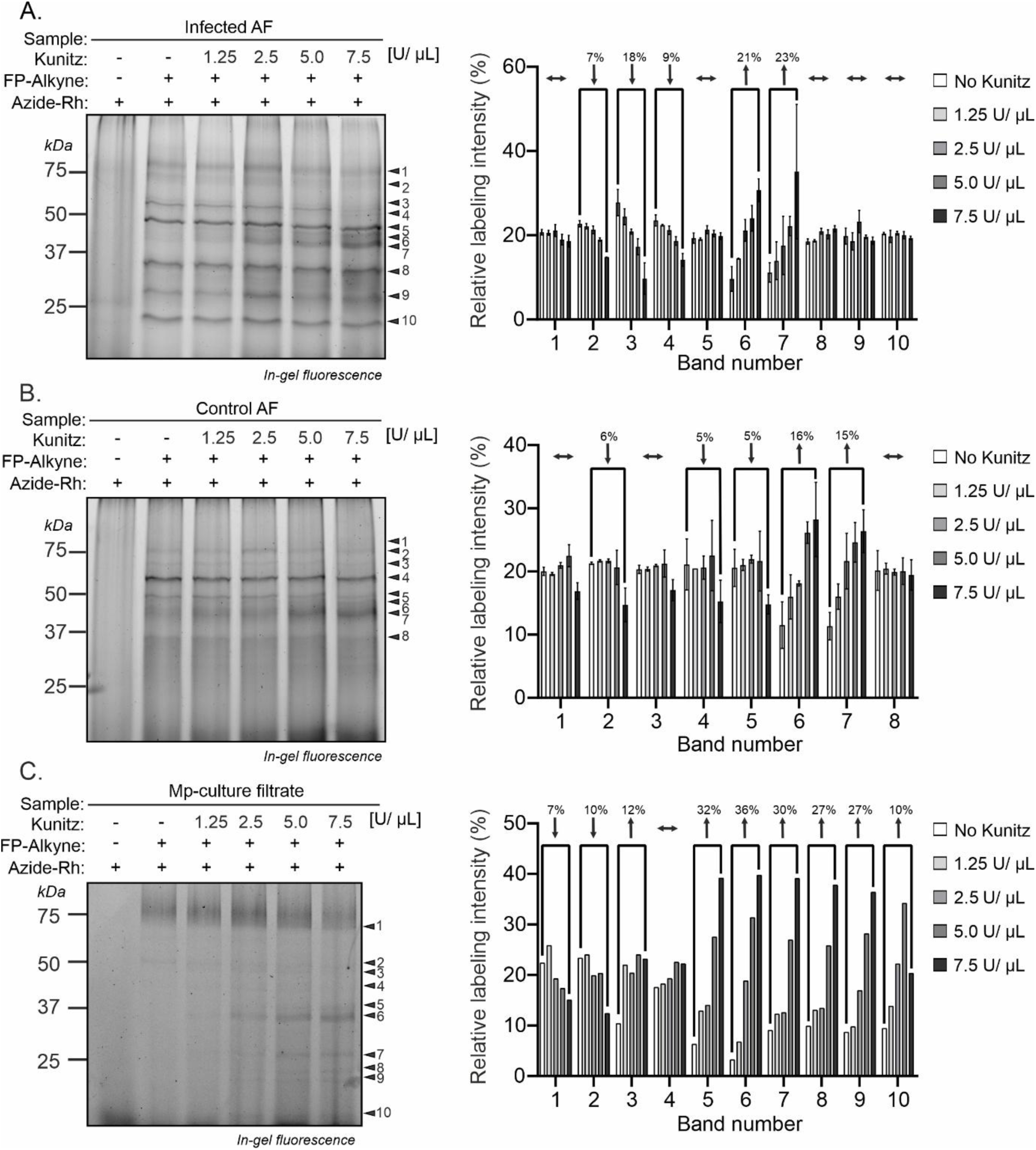
Competitive Activity-based Protein profiling (ABPP) reveals the interaction of soybean Kunitz with serine proteases of soybean and *M. phaseolina* at their active site. (A) Competitive ABPP reveals the suppression of FP-labeled signals in the *M. phaseolina* infected apoplastic fluids preincubated with commercial soybean Kunitz. (B) Competitive ABPP reveals the suppression of FP-labeled signals in control apoplastic fluids preincubated with commercial soybean Kunitz. (C) Competitive ABPP reveals the suppression of FP-labeled signals in the culture filtrates of *M. phaseolina*. The apoplastic fluids in Figure 7A and Figure 7B were isolated from soybean roots at 3 days post-infection (dpi). The culture filtrate in Figure 7C was collected from the 3-day-old liquid cultures inoculated with *M. phaseolina*. The apoplastic fluids or the culture filtrate samples were pre-incubated with various concentrations of commercially available soybean Kunitz (1.25, 2.5, 5, 7.5 U/ µL) for 30 minutes. The preincubated proteome was treated with the FP-alkyne probe for labelling and subsequently subjected to Cu-catalysed click chemistry reaction with Azide Flour 488. The fluorescently-labeled proteins were separated in 10% SDS PAGE gels under 200 v for 45 min and were detected by in-gel fluorescence using ChemiDoc MP (Biorad) scanner 532/28 excitation/emission filter of Pro-Q Emerald 488 in the blue epi filter. The arrow indicates the bands detected and are annotated accordingly. Densitometric analysis of the annotated bands was performed using Image Lab 6.1. software and the relative labeling intensity of each band was calculated and represented as a percentage. The percentage decrease or increase of labeling intensities respective to the untreated sample is indicated with the arrow mark. AF, apoplastic fluids, Mp, *M. phaseolina*, kDa, Kilo Dalton, FP, fluorophosphonate, Rh, Rhodamine.

## Discussion

*Macrophomina phaseolina* is a climate change-relevant fungal pathogen that causes charcoal root rot in various legumes, including soybean. *M. phaseolina* displays a hemibiotrophic lifestyle by employing sequential biotrophic followed by necrotrophic infection strategies to colonize their hosts (Chowdhury et al. 2017). To detect the wide array of effectors secreted by *M. phaseolina* and identify the candidate upregulated plant susceptibility factors, we have chosen the highly susceptible soybean variety, JS 20-94. In our apoplast proteomics study, we detected a large repertoire of CAZymes secreted by *M. phaseolina* in the soybean root apoplast. CAZymes are employed by fungal pathogens to degrade plant cell walls during colonization. The specific members of CAZymes secreted by *M. phaseolina* strongly correlate with the decrease in the characteristic compositions in the plant cell walls during infections. For instance, the decrease in the relative abundance of neutral sugars such as arabinose and xylose and 5-Ara*f* and 2-Ara*f* residues in the cell walls of infected soybean roots might be due to the activities of the members from the GH43 family that have been predominantly shown to exhibit α-arabinofuranosidase and/or β-xylosidase activities (Shrivastava and Goyal 2024). Reduced relative abundance of other glycosyl residues 2-Rha*p*, 4-Gal*p*, and 3,6 Gal*p* in the infected cell walls could be attributed to the activities of the members of the GH78, GH35, and GH53 families that are shown to act on these glycosyl linkages, respectively (Hobson and Deyholos 2013; Nghi et al. 2012; Tanthanuch et al. 2008). Notably, we have detected LPMOs belonging to the AA9 family secreted by *M. phaseolina* in our apoplast proteome dataset. These enzymes were not detected in the recent *M. phaseolina* secretome studies (Arafat et al. 2022; Pineda-Fretez et al. 2023). Fungal LPMOs are emerging as important virulence factors, and thus, the detection of LPMOs from *M. phaseolina* signifies the importance of our apoplastic proteome analysis. The oxidative activity of these LPMOs by *M. phaseolina* could have reduced crystalline cellulose content in the cell walls of infected soybean root. The glycosidic sugar residues that showed reduced relative abundance in the cell walls of infected samples are characteristic linkages derived from cell wall polysaccharides such as RG-1 and AGs. RG-1 and AGs exist as a complex in the plant cell walls and provide structural support to the plant cell (Tan et al. 2013). Hence, it could be speculated that *M. phaseolina* targets these cell wall components to gain entry into the soybean roots during colonization. Callose, β-1,3-glucan polymer, deposited at pathogen penetration sites helps reinforce weak or compromised cell walls during a pathogen attack (Y. Wang et al. 2021). In our study, we observed a drastic reduction in callose depositions in the infected soybean roots at different stages of *M. phaseolina* infection. The reduction in callose depositions could be due to endo-β-1,3-glucanases activities. The upregulation of plant endo-β-1,3-glucanases has resulted in callose degradation to promote viral infection and bacterial infections (Gaudioso-Pedraza et al. 2018; Shi et al. 2020). In many cases, β-1,3-glucanases are also secreted by pathogenic fungi such as *Colletotrichum graminicola* to degrade callose to promote infection (Gu et al. 2024). In our proteome dataset, we have detected various members of glycosyl hydrolases that are secreted by *M. phaseolina* (GH17, GH55 and GH131) and soybean (GH17 and GH152) that display endo-β-1,3-glucanases activity. It would be interesting to identify the candidate enzyme involved in callose degradation during infection. Taken together, it is evident that specific cell wall components are altered during infections, and hence, engineering soybean with improved resilient characteristics of these cell wall components might be an interesting approach to prevent the invasion of *M. phaseolina*.

In our proteome dataset, we have detected several candidate apoplastic effectors secreted by *M. phaseolina*. Fungal pathogens employ apoplastic effectors to promote host colonization. SUSS effectors are emerging as major players in the apoplast during plant colonization (Seong and Krasileva 2023). In our study, AF2 analysis has revealed interesting SUSS effectors secreted by *M. phaseolina*. First, the Tsp1-like effector (UniProt ID: K2RZ98) with the characteristic β-barrel structural fold. Tsp1, an effector protein, is secreted by *Trichoderma virens,* a beneficial fungus, to module host hormonal pathways, especially inducing salicylic acid in maize plants. Salicylic acid, an antagonist for Jasmonic acid, promoted susceptibility against *Cochliobolus heterostrophus*, a necrotrophic fungal pathogen (Gupta et al. 2021). Second, the effectors with KP6-like fold (UniProt ID: K2RDS7). The effectors with KP6-like fold are secreted by *Z. tritici*, a wheat pathogen, and have been reported to suppress PTI responses induced by flg22, laminarin, and chitin during the early stages of wheat colonization (Thynne et al. 2024). In addition, Zt-KP6-1, an effector with a KP6-like fold, has also been shown to function as an antimicrobial protein against fungal pathogens such as *Botrytis cinerea* and *Z. tritici* (Guillen et al. 2024). Third, MoERS1-like effectors (Uniprot IDs: K2RZH1, K2R1W5, and K2QS10) with the characteristic β-trefoil structural fold. MoERS1 effectors have been reported to be the species-specific effectors secreted by *Magnaporthe oryzae* to inhibit the papain-like cysteine protease *Os*RD21 involved in rice immunity (M. Liu et al. 2024). However, the detection of MoERS1-like effectors in our study indicates that these effectors are not species-specific but seem to be expanded in other fungi. Notably, we have detected two effectors (UniProt IDs: K2R751 and K2RET3) that had no structural similarity with the experimentally resolved structures of fungal proteins in the PDB100 database. However, these effectors have structural similarities to other Ascomycetes fungi based on our search in the AlphaFold DB. Hence, these effectors with novel structural folds might have important functions that require further investigation. Therefore, it is possible that these SUSS effectors secreted by *M. phaseolina* have similar functions as their characterized structural analogs to promote host colonization.

Fungi take advantage of certain plant proteins that act as susceptibility factors promoting colonization (Gorshkov and Tsers 2022). In our dataset, we have detected several upregulated proteins whose homologs have been reported to act as susceptibility factors in various plants. First, the upregulation of members of the GH38 family. We have detected three proteins from this family whose abundance is increased during infections. The members of this family have been shown to exhibit α-mannosidase activities and target N-glycans of cell wall glycoproteins for the cell wall loosening process (Ghosh et al. 2011; Méndez-Yáñez et al. 2024). Interestingly, increased activities of α-mannosidases have been detected in soybean roots during the colonization of arbuscular mycorrhizal fungi (AMF), and the activities of these enzymes on N-glycans have been speculated for promoting AMF colonization (Dettmann et al. 2005). Second, the upregulation of expansin proteins. In our dataset, we have two expansin proteins upregulated during infections. Expansins are involved in cell enlargement and cell wall loosening at the interaction points between cellulose and hemicellulose (Cosgrove 2005). The absence or downregulation of these proteins has resulted in increased resistance against necrotrophic fungal pathogens such as *Botrytis cinerea* and *Alternaria brassicicola* (Abuqamar et al. 2013; Cantu et al. 2008). Third, the upregulation of members of LRR-only proteins. We have detected three proteins from this family whose abundance is increased during infections. Recently, an apoplast localized LRR-only protein, NTCD4, in Arabidopsis, has been shown to act as a susceptibility factor. NTCD4 interacts with pathogen-derived Necrosis-and ethylene-inducing peptide 1 (Nep1)-like proteins (NLP) to trigger cell death and promote colonization of necrotrophic fungal pathogen *Botrytis cinerea* (Chen et al. 2021). Interestingly, we detected two NLP-like proteins (UniProt IDs: K2RMZ7, K2RNB9) secreted by *M. phaseolina* in soybean root apoplast (Table S3). Hence, it could be speculated that these proteins might exhibit interactions with the upregulated LRR-only proteins to induce cell death and promote *M. phaseolina* colonization. Fourth, the upregulation of a GH17 family protein. The members of this family exhibit endo β-1,3-glucanase activities. We have detected one GH17 family protein (Uniprot ID: I1LIL5) whose abundance is increased during infections. Recently, *Hv*BGLUII, a GH17 family protein, has been characterized for its role in releasing susceptible factors from the fungal extracellular polysaccharide (EPS) matrix. The endo β-1,3-glucanase activities of *Hv*BGLUII released a decasaccharide fragment, GD, that scavenges ROS and promotes fungal colonization (Chandrasekar et al. 2022). Taken together, these candidate susceptibility factors can be chosen as potential candidates for engineering CR resistance in soybean using genome editing tools.

Apoplastic proteases and their inhibitor interactions are significant determinants of the outcome of plant-fungal interactions. AFM is emerging as an important machining tool for investigating enzyme-inhibitor interactions at the plant-pathogen interface. Our AFM screening of selected secreted proteins has revealed the interaction of cysteine and serine protease with their respective protease inhibitors. The top AFM-predicted models of these protease inhibitor complexes are similar to the reported crystal structures of other protease inhibitor complexes where the loop regions of the protease inhibitor interact strongly with the active site of the proteases (Bode et al. 1988; Mehmood et al. 2024). Papain-like cysteine proteases (PLCPs) and serine proteases represent the central hub for plant immunity, and their activities are modulated by inhibitors during fungal colonization (Coculo et al. 2023; Misas-Villamil et al. 2016). In our study, a MoERS1-like effector (UniProt ID: K2QS10) secreted by *M. phaseolina* displayed significant interaction with a ubiquitous PLCP (UniProt ID: I1JTM0) present in the soybean root apoplast. Fungal and oomycetes pathogens secrete proteaceous inhibitors in the apoplast to modulate the activities of immune cysteine proteases during colonization (Misas-Villamil et al. 2016). Therefore, the MoERS1-like effector (UniProt ID: K2QS10) identified in this study might function as a PLCP inhibitor modulating soybean immune response. In addition, our AFM analysis has revealed interactions of soybean Kunitz with the serine protease of soybean and *M. phaseolina*. In our proteome analysis, Kunitz proteins are the only soybean-derived protease inhibitors upregulated during infections. Plant Kunitz proteins are predominantly characterized for their roles in disrupting the digestive functions of insects and animals by inhibiting the activities of serine proteases (Mehmood et al. 2024; Okedigba et al. 2023). However, plant Kunitz proteins also play important roles during plant-pathogen interactions. AtKTI1, a Kunitz protein from Arabidopsis, modulates cell death responses and also serves as a negative regulator for resistance against *Erwinia carotovora* subsp. *carotovora*, necrotrophic bacterial pathogen (Li, Brader, and Palva 2008). Furthermore, Kunitz proteins also module the activities of host serine protease in apoplast during fungal colonization. KPI106, a Kunitz protein from *Medicago truncatula,* is upregulated in root apoplast during colonization of AMF. The KPI106 inhibits an apoplast serine protease, SCP1, which is upregulated during AMF colonization. The *M. truncatula* interaction of tandemly upregulated Kunitz and serine proteases in *M. truncatula* control the colonization of AMF and arbuscule formation (Rech et al. 2013). In our dataset, the abundance of serine proteases and Kunitz are upregulated in soybean root apoplast during infections. Therefore, *M. phaseolina* might follow a strategy similar to AMF by modulating the immune serine protease activities through soybean Kunitz protein to facilitate their colonization. Aside from the negative roles of some members of Kunitz proteins in plant disease resistance, these proteins have also been shown to play positive roles against pathogens. In a recent study, StMLP1, a Kunitz protein from potato, enhanced resistance against *Ralstonia solanacearum*, a bacteria pathogen (Wang et al. 2023). Kunitz proteins have also been shown to act as antimicrobial effects on fungal pathogens under *in vitro* conditions (Clemente et al. 2019; da Silva et al. 2023). However, the exact mechanism by which Kunitz imparts resistance against plant pathogens is not known. Our AFM analysis has revealed significant interactions of soybean Kunitz proteins with the serine proteases secreted by *M. phaseolina*. So far, no reports are available for plant Kunitz proteins targeting fungal serine proteases. Fungal serine proteases are important virulence factors that have been shown to degrade plant immune enzymes, such as chitinases (Han et al. 2019; Jashni et al. 2015). Therefore, inhibiting the virulence-associated fungal serine proteases might be a mechanism of action to impart resistance toward the invading pathogens. Structured-guided engineering of protease-inhibitor complexes is emerging as an important strategy to engineer resistance in plants against pathogens. Here, key amino acid residues are mutated to disrupt the protease-inhibitor interaction, leading to enhanced resistance against pathogens (Schuster et al. 2024). Therefore, the protease-inhibitor complexes that we have predicted using AFM in our study might find their application in structured-guided engineering for improving resistance in soybean against *M. phaseolina*.

## Data Availability Statement

The scripts used for the analysis are available on github (https://github.com/chemicalglycobiologylab) and the data generated during the analysis is available in zonedo (https://zenodo.org/records/13779899). The mass spectrometry proteomics data have been deposited to the ProteomeXchange Consortium via the PRIDE [1] partner repository with the dataset identifier PXD059244. The data that support the findings of this study are available from the corresponding author upon request.

## Acknowledgments

This research work was supported by SRG grant (SRG/2022/000528) provided by Science and Engineering Research Board (SERB) India and ACRG grant provided by the Birla Institute of Technology and Science, Pilani, (BITS Pilani) Pilani campus. We thank the Central Instrumentation Facility (CIF), BITS Pilani, for providing the GC-MS/MS and confocal microscopy facilities for the research work. We thank the High-Performance Computing (HPC) Facilities, BITS Pilani for MM/PBSA simulation analysis. We also thank the Department of Biological Sciences, BITS Pilani for providing the epifluorescence microscopy and other infrastructural facilities for conducting this research work. We also thank Dr. Siddesh S Kamat and Ojal Saharan, Indian Institute of Science Education and Research (IISER), Pune, for graciously providing the FP-alkyne probe for our ABPP experiments. We thank Dr. B S Gill, Punjab Agricultural University, Ludhiana for providing the soybean seeds. We thank Department of Genetics and Plant Breeding, Jawaharlal Nehru Krishi Vishwa Vidyalaya, Jabalpur for providing the soybean seeds and the *M. phaseolina*-infected root samples. We thank BITS Pilani, Pilani Campus, for providing Institute Fellowship to Chetan Veeraganti Naveen Prakash and Muthusaravanan Sivaramakrishnan.

## Author contributions

BC, CVNP and MS conceived the project; CVNP performed pathogen isolation, infection assays, apoplastic proteome analysis, glycome analysis and competitive ABPP with the help of SG and VP; MS performed AlphaFold and AlphaFold Multimer analysis, MD simulations, callose quantification and deposition assays and Pfam analysis. PKA and MKS provided the seed material, and the *M. phaseolina*-infected root samples; BC supervised the project, designed the experiment, and wrote the manuscript with the contribution from CVNP and MS.

