## Supplementary Figures S1-S16 for "Proteome Analysis of Soybean Root Apoplast Combined with AlphaFold Prediction Reveal *Macrophomina phaseolina* Infection Strategies and Potential Targets for Engineering Resistance"

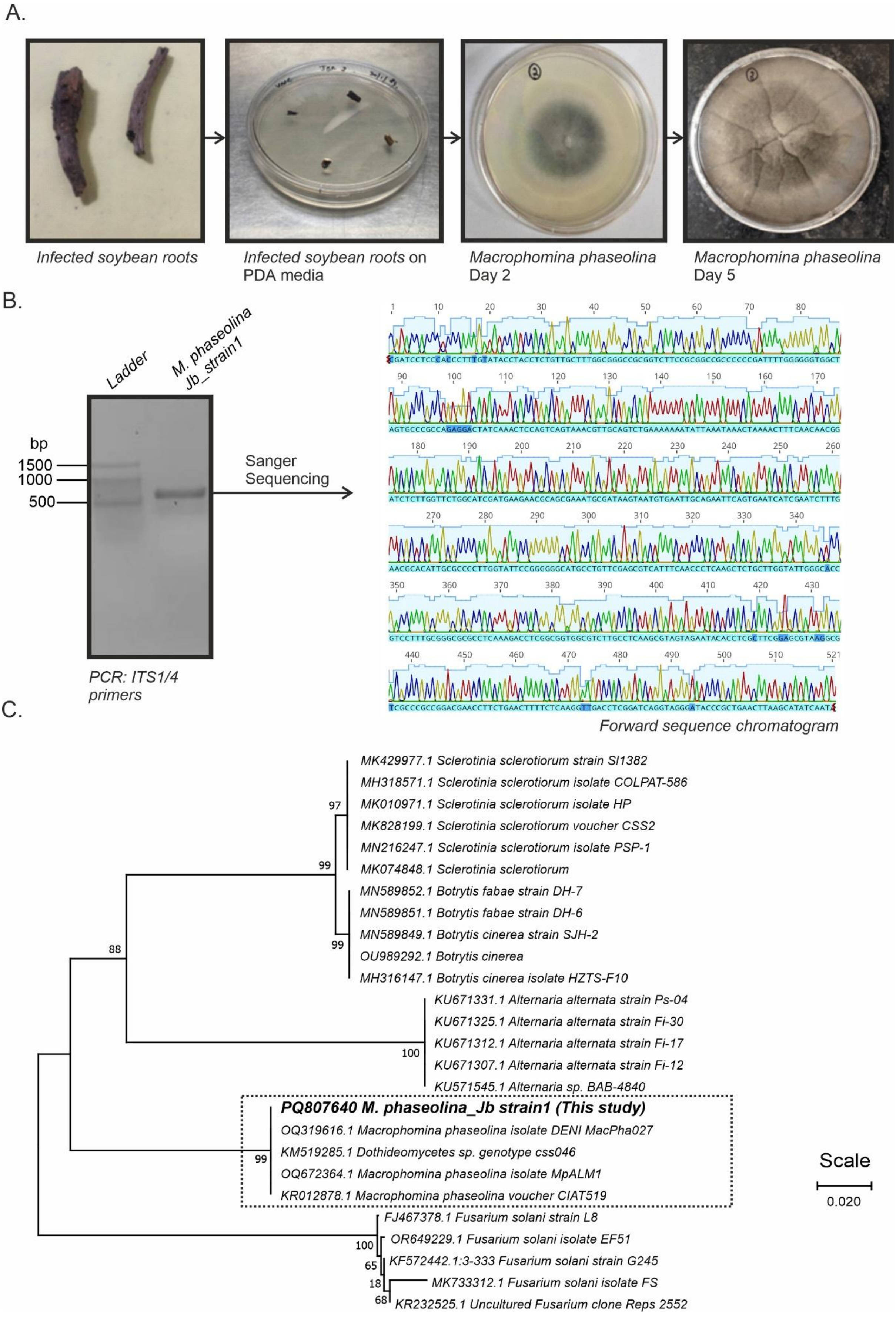

**Figure S1**: Isolation and molecular identification of *Macrophomina phaseolina* (A) Isolation of *M. phaseolina* from infected soybean roots. The infected soybean roots were collected from the fields of Jabalpur, Madhya Pradesh, India and pure culture of *M. phaseolina* was isolated on potato dextrose agar using hyphal tip method (B) Molecular identification of the isolated *M. phaseolina* strain. The fungal DNA was extracted from the pure cultures and Internal Transcribed Spacer (ITS1/ITS4) region was amplified using PCR (Primers= ITS1: TCCGTAGGTGAACCTGCGG and ITS4: TCCTCCGCTTATTGATATGC). The PCR product was resolved in an agarose gel (1.5%) using a gel electrophoresis unit and was sequenced using Sanger's method of sequencing. (C) Phylogenetic tree analysis of the *M. phaseolina_Jbstrain1*. The obtained sequence along with ITS1/ITS4 sequences of other strains of *M. phaseolina* and a few other pathogens collected from the National Center for Biotechnology Information (NCBI) database were aligned using ClustalW. The phylogenetic tree of the aligned sequences was constructed using the neighbor-joining method in MEGA 11 software (Bootstrap value= 1000, scale= 0.02). The scale denotes the genetic distance between the strains. The dotted box denotes that the *M. phaseolina_Jbstrain1* displayed high relatedness to the reported *M. phaseolina* strains. PDA, potato dextrose agar, bp, base pair.

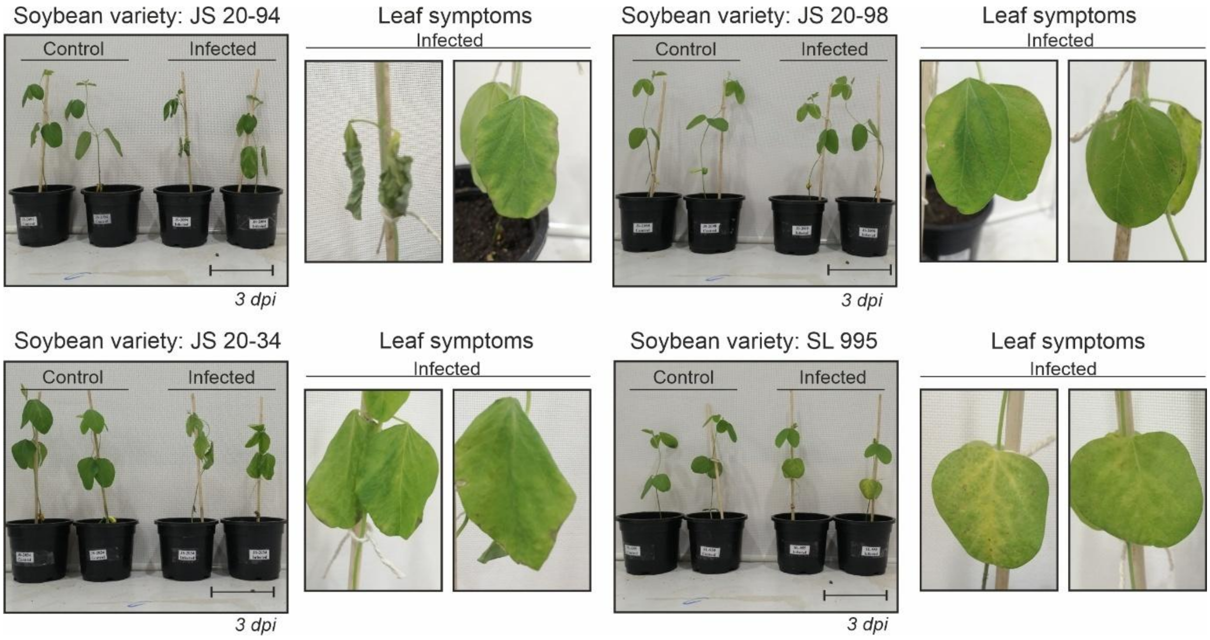


**Figure S2**: Pathogenicity test of *M. phaseolina_Jbstrain1* with elite Indian soybean varieties. The two-week-old seedlings of elite soybean varieties, JS 20-94, JS 20-34, JS 20-98 and SL 955 were treated with 25% (w/v) of homogenised actively growing *M. phaseolina* culture using root-dip inoculation method. The disease progression was observed for three days and the visual disease symptoms were recorded (Scale = 10cm). dpi, days post-infection.

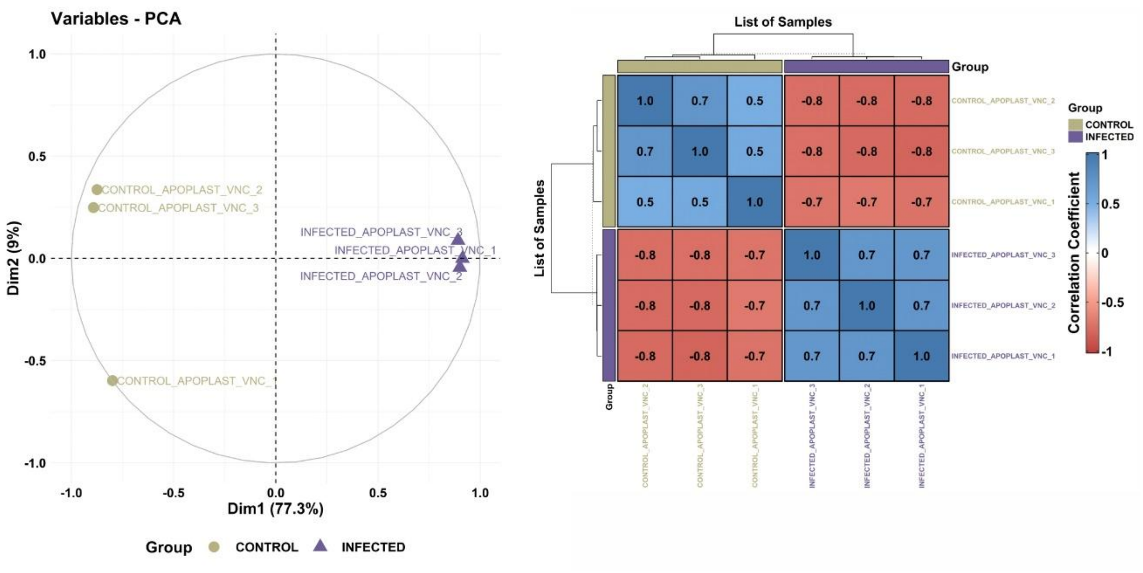


**Figure S3**: Principal Component Analysis (PCA) and correlation analysis of total apoplastic proteins. PCA and correlation was performed to show the distinctness of the control and infected apoplastic fluids and similarity within their respective biological replicates. The PCA plot was plotted using factoextra package and Pearson's correlation plot was plotted using corrplot package in R software. Dim, Dimension, PCA, Principal Component Analysis.

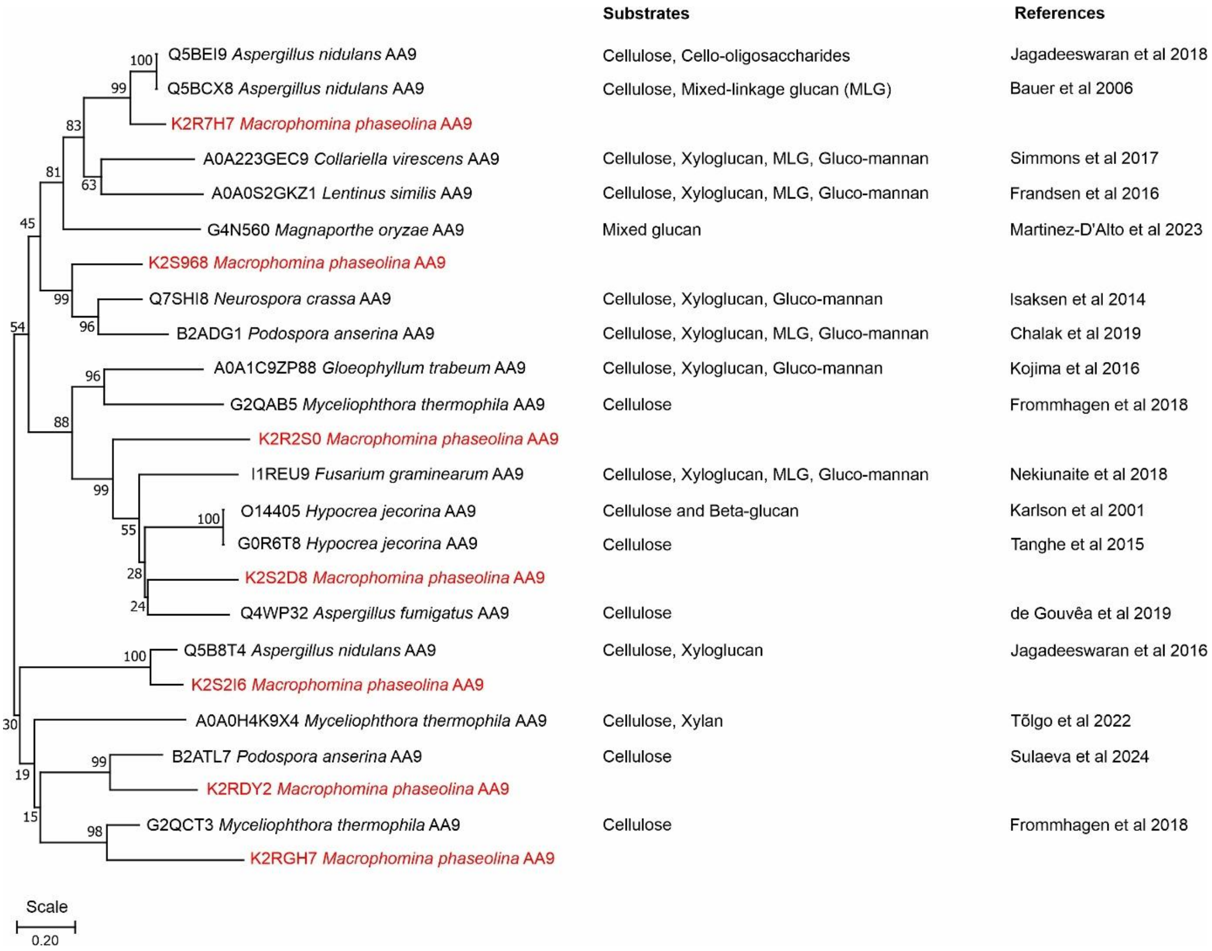


**Figure S4:** Phylogenetic tree analysis of the detected members of AA9 secreted by *M. phaseolina*. The sequences of the functionally characterized Lytic Polysaccharide Monooxygeneses (LPMOs) belonging to AA9 family of various fungi were collected from UniProt database. The collected sequences along with detected AA9 members were aligned using ClustalW and the phylogenetic tree was constructed using the neighbor-joining method in MEGA 11 software (Bootstrap value= 1000, scale= 0.02). The scale denotes the genetic distance between the members of AA9. CAZymes of the AA9 family secreted by *M. phaseolina* are denoted in red colour. The tested substrates of the reported LPMOS are listed along with their references. MLG, Mixed linkage glucans

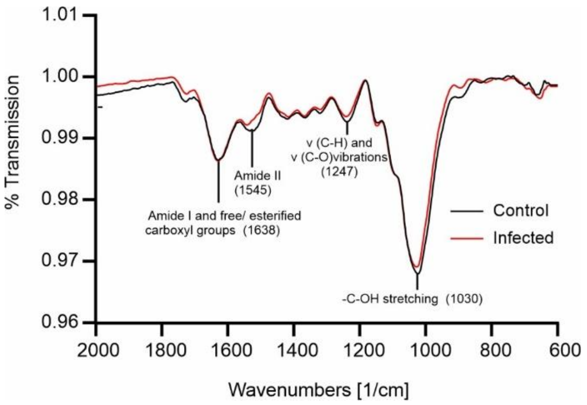


**Figure S5**: FTIR-ATR spectra analysis reveals no extensive degradation of soybean root cell wall at 3 days post-infection (dpi). FTIR analysis was performed using Fourier Transform Infrared Spectroscopy (Model: Bruker Alpha) over a range of 4,000 to 400 $cm^{-1}$. For each spectrum, 32 scans were performed at a resolution of 4 $cm^{-1}$. The spectra were baseline-corrected and smoothened using OPUS spectral processing software with default parameters. The functional groups were annotated based on corresponding wavenumbers. The sharp peak at 1638 $cm^{-1}$ corresponds to C=O group stretching vibration (Amide I) of the free and esterified carboxyl groups of pectin and proteins. The sharp peaks at 1545 $cm^{-1}$ correspond to the bending vibration of N–H and the C–N stretching vibration of the protein backbone (Amide II). The peak at 1247 and 1030 corresponds to v (C-H) and v (C-O) vibrations and -C-OH stretching of cell wall polysaccharides, respectively.

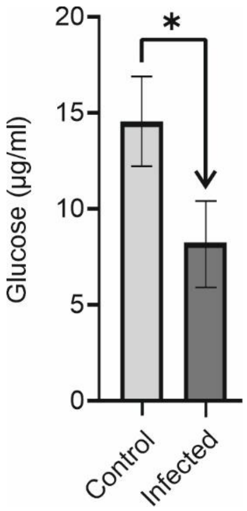


**Figure S6**: Crystalline cellulose content in the soybean root cell wall is reduced during *M. phaseolina* infection. Crystalline cellulose content was determined on the unhydrolysed pellet after the TFA hydrolysis. This pellet was treated with Updegraff reagent (acetic acid:nitric acid:water, 8:1:2 v/v) followed by Saeman hydrolysis using 72% sulphuric acid. The samples were subjected to anthrone assay to estimate the released glucose using a microtiter plate reader at 625nm. Absolute quantification of the crystalline cellulose was calculated against a standard curve plotted using glucose standards from 0-10 µg/ml. The bar represents the standard deviation between the samples. Statistical significance was determined using two-tailed t-test (*: p-value ≤ 0.05).

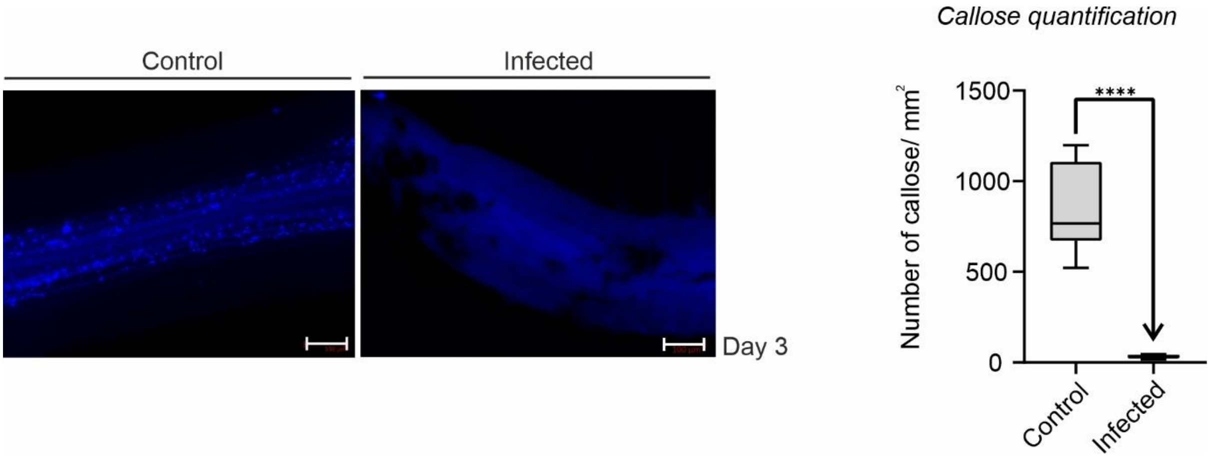


**Figure S7:** The callose depositions are reduced in soybean roots during *M. phaseolina* infection at 3 days post-infection (dpi). Aniline Blue staining was performed on the soybean roots collected at 3 dpi to visualise callose depositions. The stained lateral roots were observed under an epifluorescence microscope using a DAPI filter with an excitation wavelength of 370 nm and an emission wavelength of 509 nm (magnification= 10X, scale =100µm). Callose depositions were quantified using Trainable Weka segmentation v3.3.4 plugin of ImageJ 1.54f. The experiment was conducted with six replicates, with two replicates per plant in a completely randomized experimental design. Statistical significance was determined using two-tailed t-test (****: p-value < 0.0001).

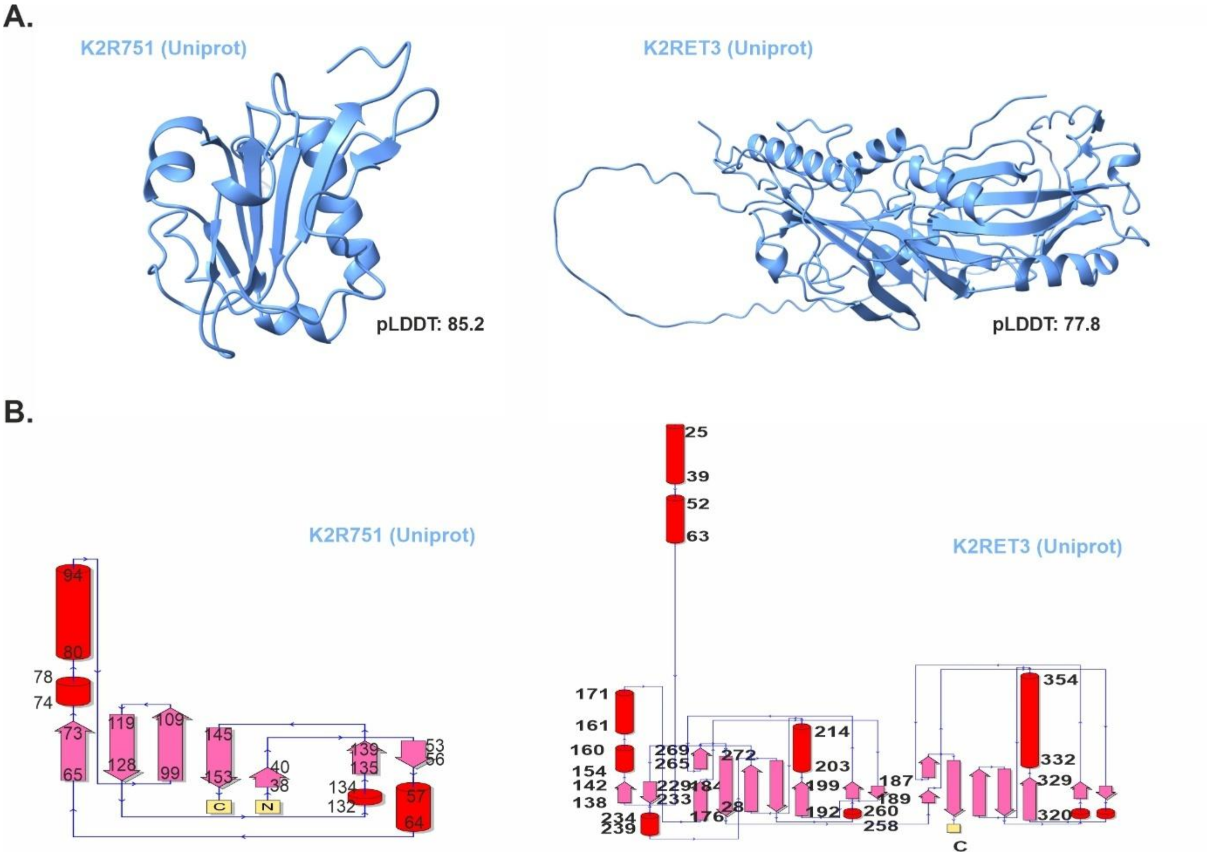


**Figure S8**: AF2-modelled structures and 2D topology maps of proteins with novel folds. (A) AF2 modelled structures of proteins K2R751 and K2RET3 with no significant similar protein hits in the PDB100 database by Foldseek algorithm. (B) The 2D topology map of proteins K2R751 and K2RET3. The 2D topology map was predicted using PDBsum webserver. pLDDT, Predicted local distance difference test score.

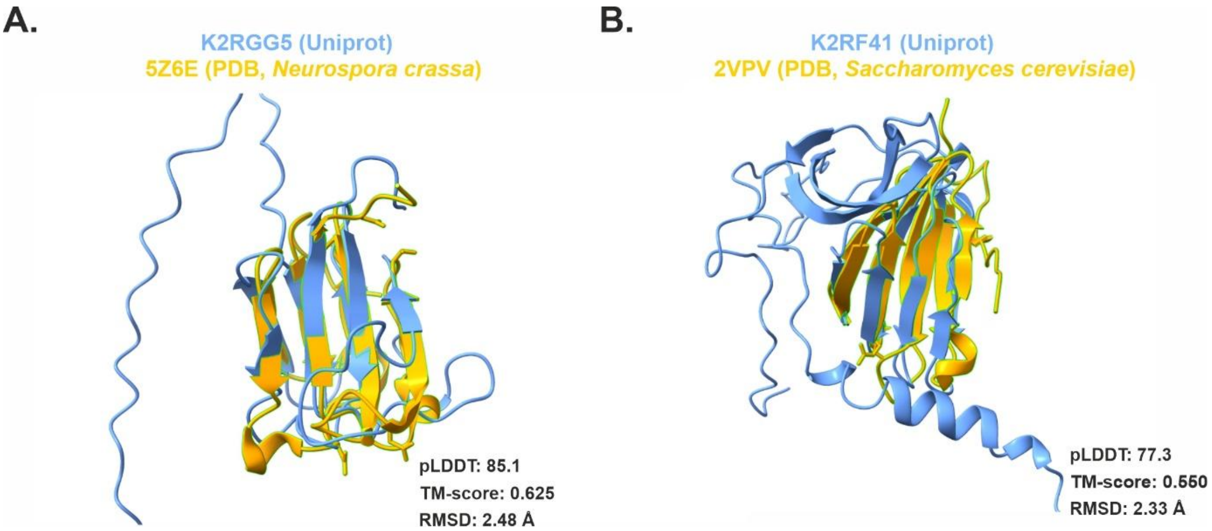


**Figure S9**: Structural similarity of AF2-modelled effectors, K2RGG5 and K2RF41 with experimentally solved crystal structures. (A) AF2-modelled effector protein K2RGG5 showing similarity to the crystal structure of the beta gamma-crystallin domain of the Abundant Perithecial Protein (APP, PDB ID: 5Z6E) from *Neurospora crassa*. (B) AF2-modelled effector protein K2RF41 demonstrating similarity to the dimerization domain of Mif2p (PDB ID: 2VPV) from *Saccharomyces cerevisiae*. The AF2 modelled proteins were superposed with structurally similar experimentally solved crystal structures available in PDB100 database using Foldseek algorithm and Matchmaker module of Chimera molecular viewer. PDB, Protein data bank, TM-score, template modelling scores, pLDDT, predicted local distance difference test score, RMSD, root mediated square deviation.

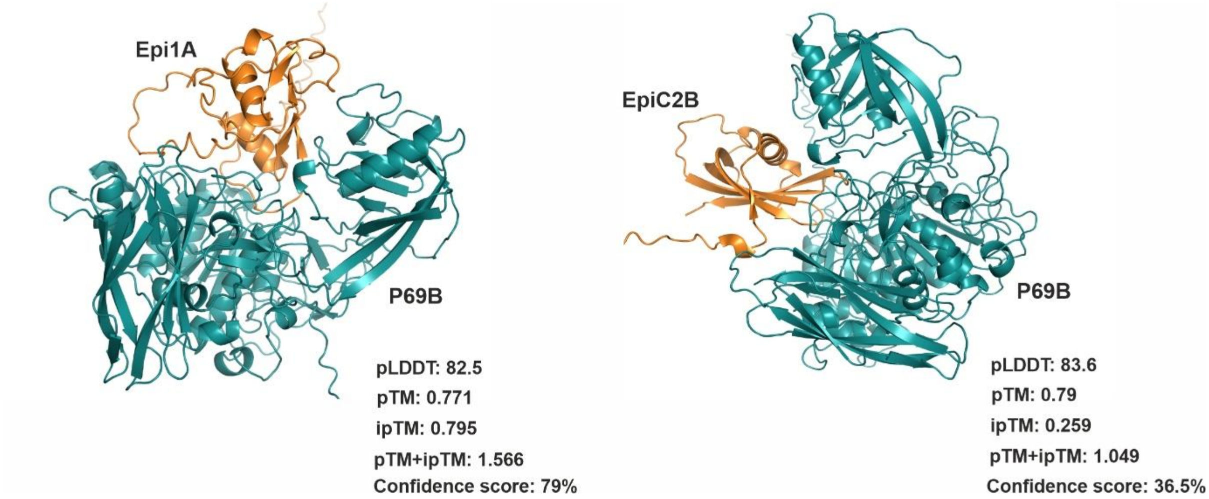


**Figure S10:** AFM predicted complexes of experimentally verified tomato serine protease-inhibitor complex Epi1a-P69B showing high ipTM value whereas swapping the inhibitor with non-inhibitor results in decreased ipTM score. This suggests that AFM can distinguish existing from non-existing complexes. pTM, predicted template modelling scores, ipTM, interface predicted template modelling scores, pLDDT, Predicted local distance difference test score.

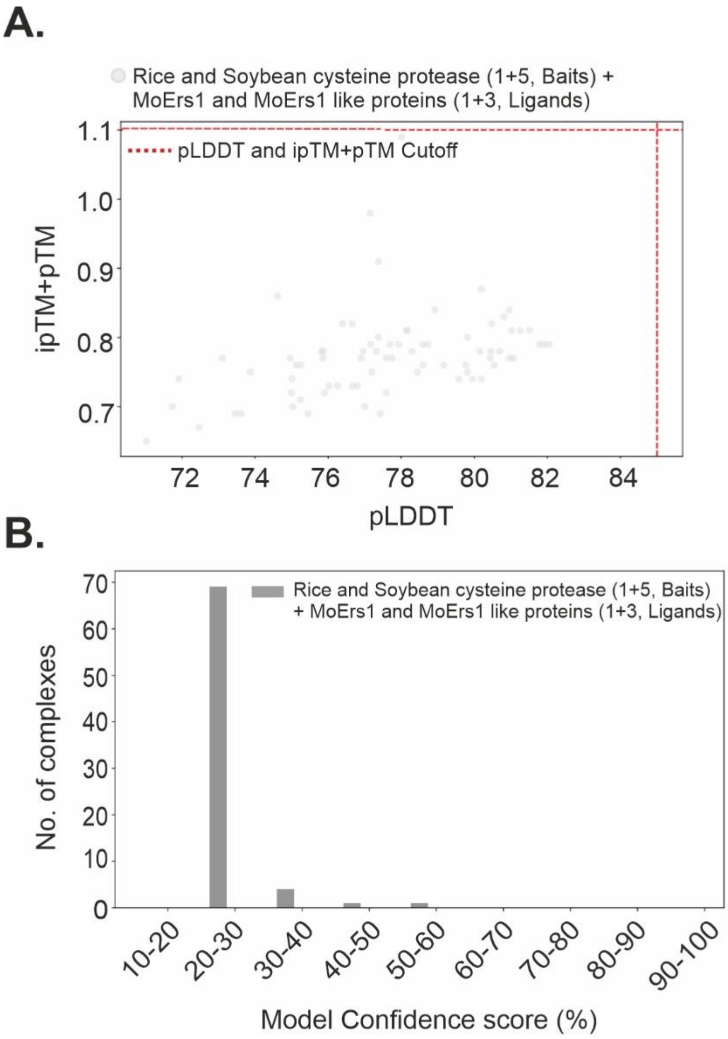


**Figure S11**: The pro-domain prevents the interaction of MoErs1 and MoErs1-like proteins with cysteine protease. AFM modelling reveals pro-domain acts as self-inhibitory region and prevents the cysteine protease active site interaction. The cysteine proteases detected in apoplasts at 3 dpi from soybean and rice RD21 protease (OsRD21) containing Pro-domain were screened against the MoErs1 effector from *Magnaporthe oryzae* and the candidate MoErs1 fold-like SUSS effector secreted by *M. phaseolina* using AlphaFold Multimer (AFM) implemented in the Localcolabfold pipeline. The performance metrics of the AFM-modelled PPI pairs are spread across the entire pTM+ ipTM score and pLDDT range, no AFM modelled PPI pairs were identified as a subpopulation of high confidence predictions with pTM+ ipTM score > 1.1 and pLDDT > 85 (upper right corner). Evaluation of the confidence score distribution (pTM*0.2 + ipTM*0.8) of AFM-predicted PPIs shows that no pairs have a confidence score > 60%, indicating that it is necessary to remove the pro-domain from the cysteine proteases before AFM modelling, as pro-domain acts as self-inhibitory region prevent the cysteine protease active site interaction with putative inhibitors. pTM, predicted template modelling scores, ipTM, interface predicted template modelling scores, pLDDT, predicted local distance difference test score.

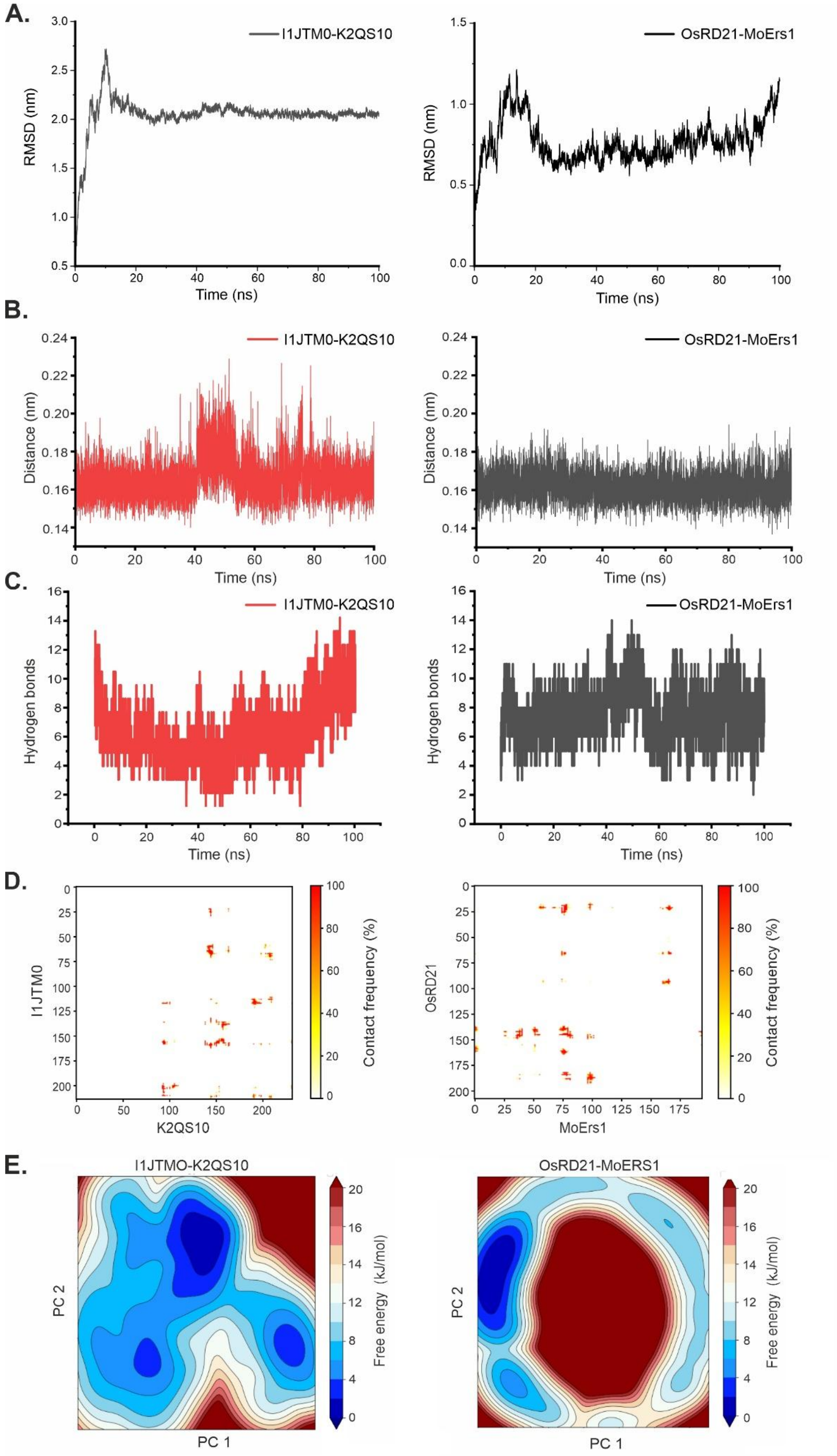


**Figure S12.** MD simulations suggest MoErs1 and MoErs1-like proteins target soybean and rice cysteine protease. (A) Root mean square deviation (RMSD) curve of the I1JTM0-K2QS10 and OsRD21-MoErs1 throughout the MD simulation of 100 ns. (B) The inter-chain distance between complexes I1JTM0-K2QS10 and OsRD21-MoErs1 during the MD simulation of 100 ns. (C) The number of hydrogen bonds at interface between complexes I1JTM0-K2QS10 and OsRD21-MoErs1 during 100 ns MD simulation. (D) The 2D contact maps show the contact frequencies for complexes I1JTM0-K2QS10 and OsRD21-MoErs1 throughout the MD simulation of 100 ns. (E) Free energy landscape showing the lowest energy conformer of the simulation trajectories for complexes OsRD21-MoErs1 and I1JTM0-K2QS10 extracted from 100 ns MD simulation. ns, nano seconds, nm, nano metre, PC, Principal Component.

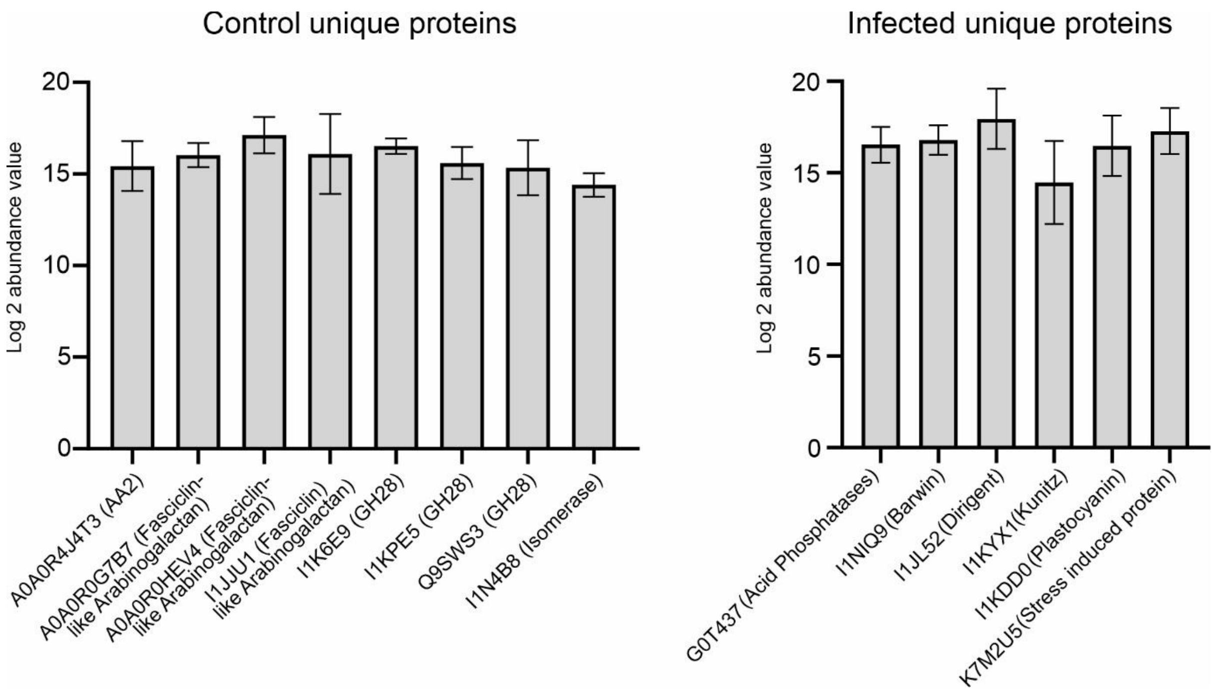


**Figure S13**: The unique soybean proteins detected in control and *M. phaseolina* infected apoplastic fluids are represented with the $log_2$ abundance values with their Uniprot ID. and domain information. The error bar represents the standard deviation within the three biological replicates.

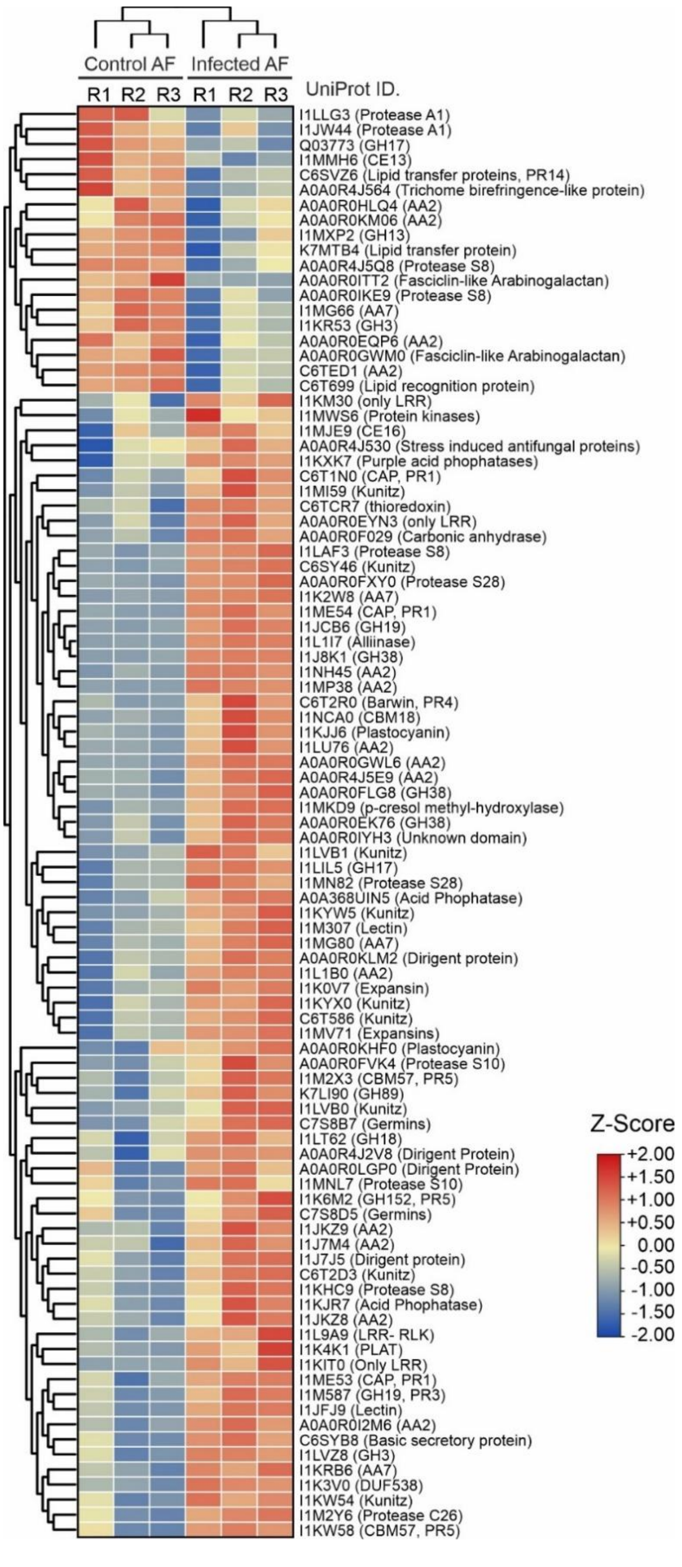


**Figure S14**: Differential expression of soybean apoplastic proteins during *M. phaseolina* infection. Heatmaps of differentially regulated proteins are plotted with z-score values using TBtools-II v1.112. Z-scores are calculated using the formula z= (X-μ)/σ, where X= individual abundance value (LFQ), μ= mean of the replicate abundance values (LFQ), σ= Standard deviation of the replicate abundance values (LFQ). The hierarchical clustering of the proteins is based on Euclidean distance, which evaluates the closeness between their expression, and the average linkage method was used for cluster analysis. Colour scale bars represent the range of z-score (-2 to +2).

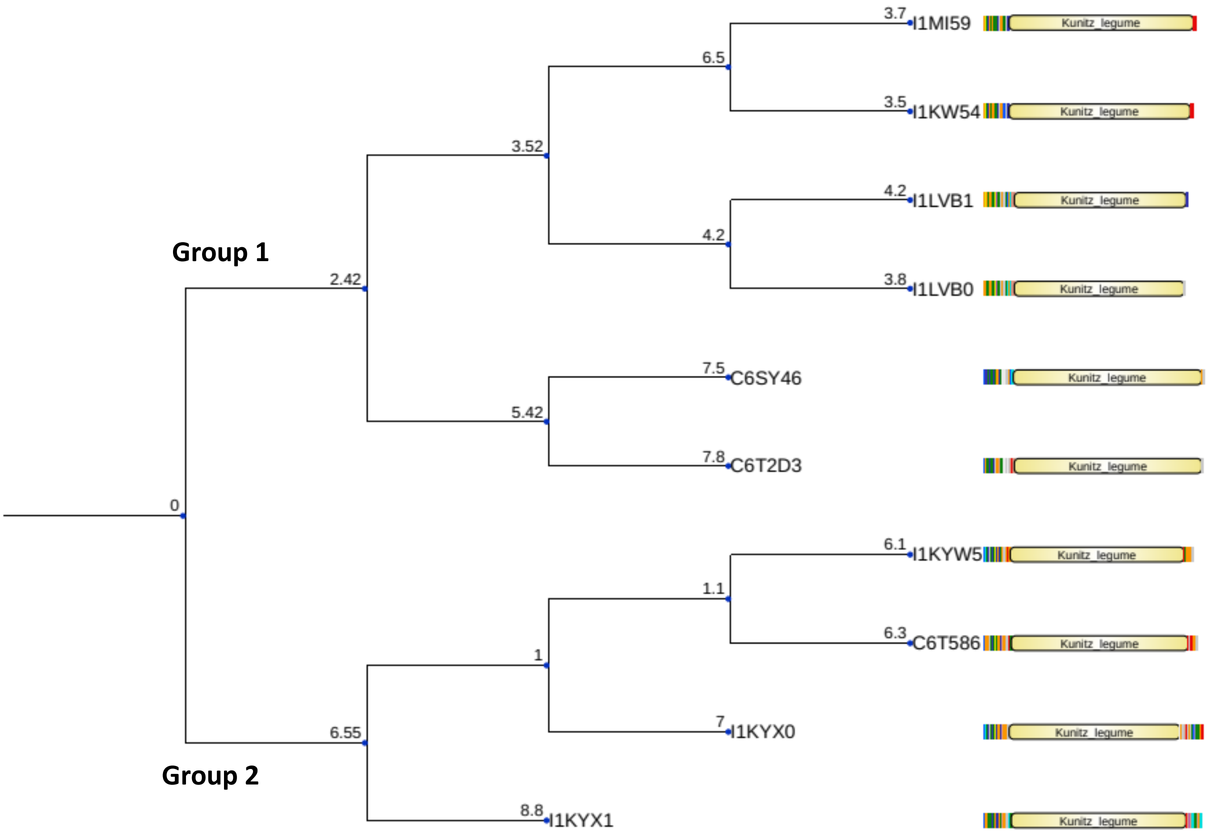


**Figure S15**: Structural comparison of soybean Kunitz proteins by DALI methodology and represented as structural similarity dendrogram. The dendrogram is derived by average linkage clustering of the structural similarity matrix (Dali Z-scores).

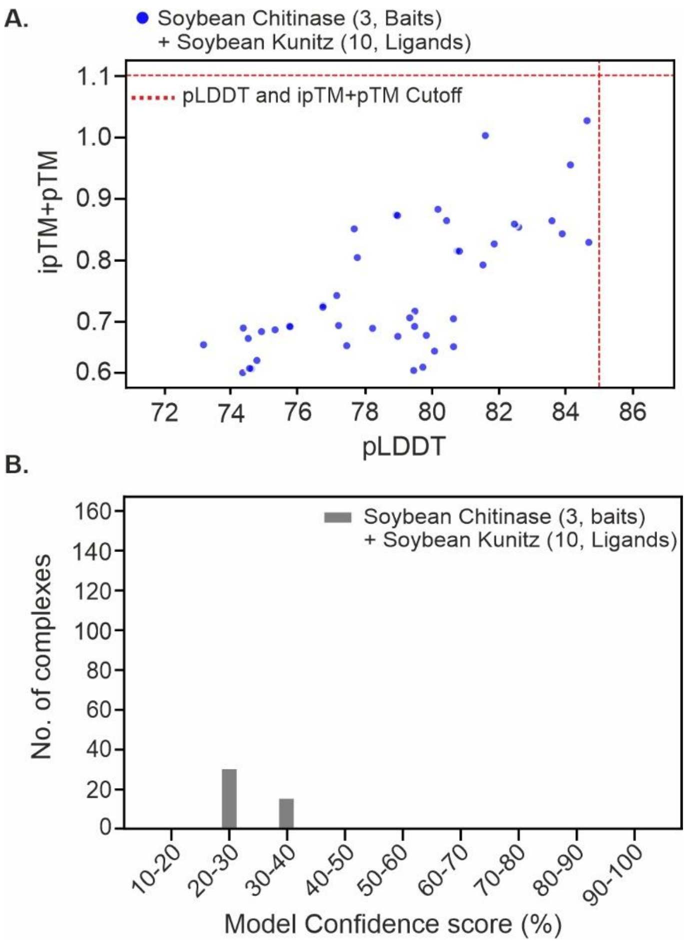


**Figure S16:** AFM reveals non-existence of interactions between soybean chitinases and soybean Kunitz. (A) The AFM predicted interaction models of soybean Kunitz and soybean chitinase distribute over the full pTM+ ipTM score and pLDDT range, with a subpopulation of highly confident predictions with pTM+ ipTM score > 1.1 and pLDDT > 85 (top right corner). The top-ranked models are coloured by dataset of origin, and highlighted in darker shade. (B) The confidence score assessment reveals no soybean Kunitz and soybean chitinase pairs potentially predicted as high confident complexes. The evaluation of confidence score distribution (pTM*0.2 + ipTM*0.8) of AFM predicted complexes, the AFM complexes with confidence score > 75% were considered as highly confident predictions.
